# Astrocytes are necessary for blood-brain barrier maintenance in the adult mouse brain

**DOI:** 10.1101/2020.03.16.993691

**Authors:** Benjamin P. Heithoff, Kijana K. George, Aubrey N. Phares, Ivan A. Zuidhoek, Carmen Munoz-Ballester, Stefanie Robel

## Abstract

In the adult brain, multiple cell types are known to produce factors that regulate blood-brain barrier properties, including astrocytes. Yet several recent studies disputed a role for mature astrocytes at the blood-brain barrier. To determine if astrocytes contribute a non-redundant and necessary function in maintaining the adult blood-brain barrier, we used a mouse model of tamoxifen-inducible astrocyte ablation. In adult mice, tamoxifen induction caused sparse apoptotic astrocyte cell death within 2 hours. Indicative of BBB damage, leakage of the small molecule Cadaverine and the large plasma protein fibrinogen into the brain parenchyma indicative of BBB damage was detected as early as astrocyte ablation was present. Vessels within and close to regions of astrocyte loss had lower expression of the tight junction protein zonula occludens-1 while endothelial glucose transporter 1 expression was undisturbed. Cadaverine leakage persisted for several weeks suggesting a lack of barrier repair. This is consistent with the finding that ablated astrocytes were not replaced. Adjacent astrocytes responded with partial non-proliferative astrogliosis, characterized by morphological changes and delayed phosphorylation of STAT3, which restricted dye leakage to the brain and vessel surface areas lacking coverage by astrocytes one month after ablation. In conclusion, astrocytes are necessary to maintain blood-brain barrier integrity in the adult brain. Blood-brain barrier-regulating factors secreted by other cell types, such as pericytes, are not sufficient to compensate for astrocyte loss.

**Main Points:** Mature astrocytes are necessary for maintenance of endothelial tight junctions in the adult brain. Ablated astrocytes are not replaced by proliferation or process extension of neighboring astrocytes resulting in long-term blood-brain barrier damage.

## 1. Introduction

A role for astrocytes in maintaining the blood-brain barrier (BBB) in the adult healthy brain is widely assumed, but little direct experimental evidence supports this conclusion. In fact, several recent studies ablating astrocytes suggest that astrocytes are not required to maintain BBB integrity (Kubotera et al., 2019; Schreiner et al., 2015; Tsai et al., 2012). The conclusion that astrocytes are involved in BBB initiation and maintenance is based on early work that transplanted glial progenitor cells into the developing main body cavity and the anterior chamber of the eye, which both normally lack a BBB. As a result of this transplantation, permeable vessels acquired BBB properties and dyes such as trypan blue and Evans blue could no longer pass this barrier (Janzer & Raff, 1987; Stewart & Wiley, 1981). Evidence for an astrocyte-secreted factor responsible for initiating barrier-like properties in endothelial cells was obtained in culture studies using primary bovine brain endothelial cells in conjunction with primary postnatal astrocytes or their conditioned media. Under these conditions, endothelial cells that lose their BBB properties when cultured alone start to express higher levels of tight junction proteins, and enhanced trans-endothelial electrical resistance (TEER) can be measured suggesting a physically tighter barrier (Rubin et al., 1991; Tao-Cheng, Nagy, & Brightman, 1987; Wolburg et al., 1994). These results were reproduced in human brain microvascular endothelial cells (HBMECs) using conditioned media from human astrocytes (Siddharthan, Kim, Liu, & Kim, 2007) or induced pluripotent stem cell-derived astrocytes (Canfield et al., 2017). From this data, a key role for astrocytes in initiating and maintaining the BBB via secreted factors, among them sonic hedgehog (Shh) and glial derived neurotrophic factor (Igarashi et al., 1999; Xia et al., 2013) was deduced. Yet astrocytes cultured from neonates rapidly divide and reflect a more progenitor-like phenotype, which is vastly different from the mature, quiescent astrocytes populating the adult brain. In 2010, Armulik and Daneman challenged the notion that BBB formation during development relies on astrocytes, arguing that the BBB is already established before glial progenitor cells differentiate into astrocytes and astrocyte endfeet are formed along vessels. Instead, this study established a role for pericytes in initiating BBB formation because the barrier was found to be abnormal in PDGFRb-deficient mice, which present with reduced pericyte coverage along cerebral vessels. Given the developmentally dysfunctional vessels and BBB, this study was not able to dissect a potential role for mature astrocytes in BBB maintenance upstream of pericytes in the adult brain (Armulik et al., 2010; Daneman, Zhou, Kebede, & Barres, 2010).

Many factors secreted by cultured astrocytes that were reported to modulate the BBB *in vitro,* including Sonic hedgehog (Shh), glial-derived neurotrophic factor and angiopoietin, are not present in mature astrocytes in the uninjured brain according to several astrocyte-specific transcriptome and proteome datasets (Anderson et al., 2016; Sharma et al., 2015; Y. Zhang et al., 2014). This holds true even after enriching for astrocyte endfeet transcripts (Boulay et al., 2017). A study carefully assessing Shh protein and mRNA expression patterns determined that Shh is present in very few astrocytes and the majority of Shh protein in the adult uninjured cortex is made by neurons (Sirko et al., 2013). Other factors including Fibroblast growth factor 2 (FGF2) and vascular endothelial growth factor A are expressed at low or, in the case of angiotensinogen, high levels. Many of these molecules are modulated in expression during development and after injury and their role for BBB function has mostly been explored in the context of pathology (Argaw et al., 2012; Huang et al., 2012; Lee et al., 2003; Min et al., 2015; Z. G. Zhang, Zhang, Croll, & Chopp, 2002; Z. G. Zhang et al., 2000; Zheng et al., 2009) or in genetic models that did not distinguish developmental or cell type-specific effects *in vivo* (Kakinuma et al., 1998; Reuss, Dono, & Unsicker, 2003; Wosik et al., 2007) to assess the role of mature astrocytes in BBB maintenance.

Thus, whether astrocytes have a necessary and non-redundant role in maintenance of the BBB in the healthy adult brain is still unresolved as the above-mentioned astrocyte ablation studies used readouts that only represent massive damage to the BBB (also see discussion). To overcome these limitations, we genetically ablated astrocytes in the adult brain and employed sensitive readouts to test for BBB integrity. Here, we demonstrate that astrocytes perform necessary and non-redundant roles for maintenance and repair of the BBB.

## 2. Methods

### 2.1 Mice

To enable inducible genetic astrocyte-specific ablation Gt(ROSA)26Sortm1(DTA)Jpmb/J mice (Jackson Labs stock #006331) were crossed with Tg(Slc1a3-cre/ERT)1Nat (Nathans, 2010) (Jackson Labs stock #012586). We will refer to Gt(ROSA)26Sortm1(DTA)Jpmb/J mice expressing one diphtheria toxin A (DTA) allele as DTA^fl/wt^ mice and to Tg(Slc1a3-cre/ERT)1Nat expressing the transgene heterozygously as Glast-CreERT^tg/wt^ mice.

Mice were initially purchased from The Jackson Laboratory and were then bred in-house. All animal procedures were approved and carried out according to the guidelines of the Institutional Animal Care and Use Committee of Virginia Polytechnic Institute and State University (Virginia Tech) and were performed in compliance with the National Institute of Health’s *Guide for the Care and Use of Laboratory Animals*.

### 2.2 Experimental Design

8-12 week old mice of both sexes were used for experiments. Sex is specified in **Table 1**.

**Table 1.** Sex and genotype of mice used in experiments.

| Figure 1 | Panel | Animal # | Sex | Experimental Group | Timepoint | Total n |
| --- | --- | --- | --- | --- | --- | --- |
|  | c | 0046 | M | DTA<sup>fl/wt</sup>//Glast-CreERT<sup>tg/wt</sup> + Tamoxifen | 1 dpa |  |
|  | d | 4520 | F | DTA<sup>wt/wt</sup>//Glast-CreERT<sup>tg/wt</sup> + Tamoxifen | 3 dpa |  |
|  |  | 309 | F | DTA<sup>fl/wt</sup>//Glast-CreERT<sup>tg/wt</sup> + Tamoxifen | 6 hpa |  |
|  |  | 305 | M | DTA<sup>fl/wt</sup>//Glast-CreERT<sup>tg/wt</sup> + Tamoxifen | 11 dpa |  |
|  |  | 4624 | M | DTA<sup>fl/wt</sup>//Glast-CreERT<sup>tg/wt</sup> + Tamoxifen | 28 dpa |  |
|  | e | 4864 | F | DTA<sup>fl/wt</sup>//Glast-CreERT<sup>wt/wt</sup> + Tamoxifen | 28 dpa |  |
|  |  | 315 | M | DTA<sup>fl/wt</sup>//Glast-CreERT<sup>tg/wt</sup> + Tamoxifen | 11 dpa |  |
|  | f,g | 528 | F | DTA<sup>wt/wt</sup>//Glast-CreERT<sup>tg/wt</sup> + Tamoxifen | 6 hpa | n=15 (control) |
|  |  | 532 | F | DTA<sup>wt/wt</sup>//Glast-CreERT<sup>tg/wt</sup> + Tamoxifen | 6 hpa |  |
|  |  | 319 | F | DTA<sup>wt/wt</sup>//Glast-CreERT<sup>tg/wt</sup> + Tamoxifen | 6 hpa |  |
|  |  | 4902 | M | DTA<sup>wt/wt</sup>//Glast-CreERT<sup>tg/wt</sup> + Tamoxifen | 1 dpa |  |
|  |  | 292 | F | DTA<sup>fl/wt</sup>//Glast-CreERT<sup>wt/wt</sup> + Tamoxifen | 1 dpa |  |
|  |  | 526 | F | DTA<sup>wt/wt</sup>//Glast-CreERT<sup>tg/wt</sup> + Tamoxifen | 1 dpa |  |
|  |  | 0043 | F | DTA<sup>fl/wt</sup>//Glast-CreERT<sup>wt/wt</sup> + Tamoxifen | 1 dpa |  |
|  |  | 4287 | F | DTA<sup>fl/wt</sup>//Glast-CreERT<sup>wt/wt</sup> + Tamoxifen | 1 dpa |  |
|  |  | 4520 | F | DTA<sup>fl/wt</sup>//Glast-CreERT<sup>wt/wt</sup> + Tamoxifen | 3 dpa |  |
|  |  | 3358 | M | DTA<sup>fl/wt</sup>//Glast-CreERT<sup>wt/wt</sup> + Tamoxifen | 3 dpa |  |
|  |  | 313 | M | DTA<sup>wt/wt</sup>//Glast-CreERT<sup>tg/wt</sup> + Tamoxifen | 11 dpa |  |
|  |  | 4864 | M | DTA<sup>fl/wt</sup>//Glast-CreERT<sup>wt/wt</sup> + Tamoxifen | 28 dpa |  |
|  |  | 4862 | M | DTA<sup>fl/wt</sup>//Glast-CreERT<sup>wt/wt</sup> + Tamoxifen | 28 dpa |  |
|  |  | 219 | F | DTA<sup>wt/wt</sup>//Glast-CreERT<sup>wt/wt</sup> + Tamoxifen | 28 dpa |  |
|  |  | 4995 | F | DTA<sup>fl/wt</sup>//Glast-CreERT<sup>wt/wt</sup> + Tamoxifen | 28 dpa |  |
|  |  | 893 | F | DTA<sup>fl/wt</sup>//Glast-CreERT<sup>tg/wt</sup> + Tamoxifen | 2 hpa | n=3 (2 hpa) |
|  |  | 895 | M | DTA<sup>fl/wt</sup>//Glast-CreERT<sup>tg/wt</sup> + Tamoxifen | 2 hpa |  |
|  |  | 892 | M | DTA<sup>fl/wt</sup>//Glast-CreERT<sup>tg/wt</sup> + Tamoxifen | 2 hpa |  |
|  |  | 309 | F | DTA<sup>fl/wt</sup>//Glast-CreERT<sup>tg/wt</sup> + Tamoxifen | 6 hpa | n=3 (6 hpa) |
|  |  | 868 | M | DTA<sup>fl/wt</sup>//Glast-CreERT<sup>tg/wt</sup> + Tamoxifen | 6 hpa |  |
|  |  | 723 | F | DTA<sup>fl/wt</sup>//Glast-CreERT<sup>tg/wt</sup> + Tamoxifen | 6 hpa |  |
|  |  | 4901 | M | DTA<sup>fl/wt</sup>//Glast-CreERT<sup>tg/wt</sup> + Tamoxifen | 1 dpa | n=4 (1 dpa) |
|  |  | 294 | F | DTA<sup>fl/wt</sup>//Glast-CreERT<sup>tg/wt</sup> + Tamoxifen | 1 dpa |  |
|  |  | 3497 | F | DTA<sup>fl/wt</sup>//Glast-CreERT<sup>tg/wt</sup> + Tamoxifen | 1 dpa |  |
|  |  | 4221 | F | DTA<sup>fl/wt</sup>//Glast-CreERT<sup>tg/wt</sup> + Tamoxifen | 1 dpa |  |
|  |  | 4519 | M | DTA<sup>fl/wt</sup>//Glast-CreERT<sup>tg/wt</sup> + Tamoxifen | 3 dpa | n=4 (3 dpa) |
|  |  | 4523 | M | DTA<sup>fl/wt</sup>//Glast-CreERT<sup>tg/wt</sup> + Tamoxifen | 3 dpa |  |
|  |  | 3351 | M | DTA<sup>fl/wt</sup>//Glast-CreERT<sup>tg/wt</sup> + Tamoxifen | 3 dpa |  |
|  |  | 2279 | M | DTA<sup>fl/wt</sup>//Glast-CreERT<sup>tg/wt</sup> + Tamoxifen | 3 dpa |  |
|  |  | 538 | M | DTA<sup>fl/wt</sup>//Glast-CreERT<sup>tg/wt</sup> + Tamoxifen | 5 dpa | n=4 (5 dpa) |
|  |  | 317 | M | DTA<sup>fl/wt</sup>//Glast-CreERT<sup>tg/wt</sup> + Tamoxifen | 5 dpa |  |
|  |  | 535 | F | DTA<sup>fl/wt</sup>//Glast-CreERT<sup>tg/wt</sup> + Tamoxifen | 5 dpa |  |
|  |  | 536 | F | DTA<sup>fl/wt</sup>//Glast-CreERT<sup>tg/wt</sup> + Tamoxifen | 5 dpa |  |
|  |  | 310 | M | DTA<sup>fl/wt</sup>//Glast-CreERT<sup>tg/wt</sup> + Tamoxifen | 11 dpa | n=4 (11 dpa) |
|  |  | 305 | M | DTA<sup>fl/wt</sup>//Glast-CreERT<sup>tg/wt</sup> + Tamoxifen | 11 dpa |  |
|  |  | 529 | M | DTA<sup>fl/wt</sup>//Glast-CreERT<sup>tg/wt</sup> + Tamoxifen | 11 dpa |  |
|  |  | 314 | F | DTA<sup>fl/wt</sup>//Glast-CreERT<sup>tg/wt</sup> + Tamoxifen | 11 dpa |  |
|  |  | 4624 | M | DTA<sup>fl/wt</sup>//Glast-CreERT<sup>tg/wt</sup> + Tamoxifen | 28 dpa | n=5 (28 dpa) |
|  |  | 4631 | F | DTA<sup>fl/wt</sup>//Glast-CreERT<sup>tg/wt</sup> + Tamoxifen | 28 dpa |  |
|  |  | 220 | F | DTA<sup>fl/wt</sup>//Glast-CreERT<sup>tg/wt</sup> + Tamoxifen | 28 dpa |  |
|  |  | 291 | F | DTA<sup>fl/wt</sup>//Glast-CreERT<sup>tg/wt</sup> + Tamoxifen | 28 dpa |  |
|  |  | 4836 | M | DTA<sup>fl/wt</sup>//Glast-CreERT<sup>tg/wt</sup> + Tamoxifen | 28 dpa |  |
| Figure 2 | a | 4518 | F | DTA<sup>fl/wt</sup>//Glast-CreERT<sup>wt/wt</sup> + Tamoxifen | 3 dpa |  |
|  |  | 309 | F | DTA<sup>fl/wt</sup>//Glast-CreERT<sup>tg/wt</sup> + Tamoxifen | 6 hpa |  |
|  |  | 314 | F | DTA<sup>fl/wt</sup>//Glast-CreERT<sup>tg/wt</sup> + Tamoxifen | 11 dpa |  |
|  |  | 4631 | F | DTA<sup>fl/wt</sup>//Glast-CreERT<sup>tg/wt</sup> + Tamoxifen | 28 dpa |  |
|  | b | 4902 | M | DTA<sup>wt/wt</sup>//Glast-CreERT<sup>tg/wt</sup> + Tamoxifen | 1 dpa |  |
|  |  | 305 | M | DTA<sup>fl/wt</sup>//Glast-CreERT<sup>tg/wt</sup> + Tamoxifen | 11 dpa |  |
|  | c | 319 | F | DTA<sup>wt/wt</sup>//Glast-CreERT<sup>tg/wt</sup> + Tamoxifen | 6 hpa | n=16 (control) |
|  |  | 528 | F | DTA<sup>wt/wt</sup>//Glast-CreERT<sup>tg/wt</sup> + Tamoxifen | 6 hpa |  |
|  |  | 532 | F | DTA<sup>wt/wt</sup>//Glast-CreERT<sup>tg/wt</sup> + Tamoxifen | 6 hpa |  |
|  |  | 4902 | M | DTA<sup>fl/wt</sup>//Glast-CreERT<sup>wt/wt</sup> + Tamoxifen | 1 dpa |  |
|  |  | 292 | F | DTA<sup>fl/wt</sup>//Glast-CreERT<sup>wt/wt</sup> + Tamoxifen | 1 dpa |  |
|  |  | A8032 | M | DTA<sup>fl/wt</sup>//Glast-CreERT<sup>wt/wt</sup> + Tamoxifen | 3 dpa |  |
|  |  | 4518 | F | DTA<sup>fl/wt</sup>//Glast-CreERT<sup>wt/wt</sup> + Tamoxifen | 3 dpa |  |
|  |  | 4520 | F | DTA<sup>wt/wt</sup>//Glast-CreERT<sup>tg/wt</sup> + Tamoxifen | 3 dpa |  |
|  |  | 537 | M | DTA<sup>fl/wt</sup>//Glast-CreERT<sup>wt/wt</sup> + Tamoxifen | 5 dpa |  |
|  |  | 4862 | M | DTA<sup>fl/wt</sup>//Glast-CreERT<sup>wt/wt</sup> + Tamoxifen | 28 dpa |  |
|  |  | 219 | F | DTA<sup>wt/wt</sup>//Glast-CreERT<sup>wt/wt</sup> + Tamoxifen | 28 dpa |  |
|  |  | 4629 | F | DTA<sup>fl/wt</sup>//Glast-CreERT<sup>wt/wt</sup> + Tamoxifen | 28 dpa |  |
|  |  | 272 | F | DTA<sup>fl/wt</sup>//Glast-CreERT<sup>wt/wt</sup> + Tamoxifen | 28 dpa |  |
|  |  | 273 | F | DTA<sup>fl/wt</sup>//Glast-CreERT<sup>wt/wt</sup> + Tamoxifen | 28 dpa |  |
|  |  | 4864 | M | DTA<sup>fl/wt</sup>//Glast-CreERT<sup>wt/wt</sup> + Tamoxifen | 28 dpa |  |
|  |  | 4995 | F | DTA<sup>fl/wt</sup>//Glast-CreERT<sup>wt/wt</sup> + Tamoxifen | 28 dpa |  |
|  |  | 893 | F | DTA<sup>fl/wt</sup>//Glast-CreERT<sup>tg/wt</sup> + Tamoxifen | 2 hpa | n=3 (2 hpa) |
|  |  | 895 | M | DTA<sup>fl/wt</sup>//Glast-CreERT<sup>tg/wt</sup> + Tamoxifen | 2 hpa |  |
|  |  | 892 | M | DTA<sup>fl/wt</sup>//Glast-CreERT<sup>tg/wt</sup> + Tamoxifen | 2 hpa |  |
|  |  | 309 | F | DTA<sup>fl/wt</sup>//Glast-CreERT<sup>tg/wt</sup> + Tamoxifen | 6 hpa | n=3 (6 hpa) |
|  |  | 723 | F | DTA<sup>fl/wt</sup>//Glast-CreERT<sup>tg/wt</sup> + Tamoxifen | 6 hpa |  |
|  |  | 868 | M | DTA<sup>fl/wt</sup>//Glast-CreERT<sup>tg/wt</sup> + Tamoxifen | 6 hpa |  |
|  |  | 4901 | M | DTA<sup>fl/wt</sup>//Glast-CreERT<sup>tg/wt</sup> + Tamoxifen | 1 dpa | n=3 (1 dpa) |
|  |  | 293 | F | DTA<sup>fl/wt</sup>//Glast-CreERT<sup>tg/wt</sup> + Tamoxifen | 1 dpa |  |
|  |  | 294 | F | DTA<sup>fl/wt</sup>//Glast-CreERT<sup>tg/wt</sup> + Tamoxifen | 1 dpa |  |
|  |  | 4519 | M | DTA<sup>fl/wt</sup>//Glast-CreERT<sup>tg/wt</sup> + Tamoxifen | 3 dpa | n=3 (3 dpa) |
|  |  | 4523 | M | DTA<sup>fl/wt</sup>//Glast-CreERT<sup>tg/wt</sup> + Tamoxifen | 3 dpa |  |
|  |  | A8033 | M- | DTA<sup>fl/wt</sup>//Glast-CreERT<sup>tg/wt</sup> + Tamoxifen | 3 dpa |  |
|  |  | 536 | F | DTA<sup>fl/wt</sup>//Glast-CreERT<sup>tg/wt</sup> + Tamoxifen | 5 dpa | n=3 (5 dpa) |
|  |  | 535 | F | DTA<sup>fl/wt</sup>//Glast-CreERT<sup>tg/wt</sup> + Tamoxifen | 5 dpa |  |
|  |  | 317 | M | DTA<sup>fl/wt</sup>//Glast-CreERT<sup>tg/wt</sup> + Tamoxifen | 5 dpa |  |
|  |  | 314 | F | DTA<sup>fl/wt</sup>//Glast-CreERT<sup>tg/wt</sup> + Tamoxifen | 11 dpa | n=4 (11 dpa) |
|  |  | 305 | M | DTA<sup>fl/wt</sup>//Glast-CreERT<sup>tg/wt</sup> + Tamoxifen | 11 dpa |  |
|  |  | 310 | F | DTA<sup>fl/wt</sup>//Glast-CreERT<sup>tg/wt</sup> + Tamoxifen | 11 dpa |  |
|  |  | 310 | M | DTA<sup>fl/wt</sup>//Glast-CreERT<sup>tg/wt</sup> + Tamoxifen | 11 dpa |  |
|  |  | 4631 | F | DTA<sup>fl/wt</sup>//Glast-CreERT<sup>tg/wt</sup> + Tamoxifen | 28 dpa | n=5 (28 dpa) |
|  |  | 4624 | M | DTA<sup>fl/wt</sup>//Glast-CreERT<sup>tg/wt</sup> + Tamoxifen | 28 dpa |  |
|  |  | 4836 | M | DTA<sup>fl/wt</sup>//Glast-CreERT<sup>tg/wt</sup> + Tamoxifen | 28 dpa |  |
|  |  | 291 | F | DTA<sup>fl/wt</sup>//Glast-CreERT<sup>tg/wt</sup> + Tamoxifen | 28 dpa |  |
|  |  | 220 | F | DTA<sup>fl/wt</sup>//Glast-CreERT<sup>tg/wt</sup> + Tamoxifen | 28 dpa |  |
|  | d | 4518 | F | DTA<sup>fl/wt</sup>//Glast-CreERT<sup>wt/wt</sup> + Tamoxifen | 3 dpa |  |
|  |  | 294 | F | DTA<sup>fl/wt</sup>//Glast-CreERT<sup>tg/wt</sup> + Tamoxifen | 1 dpa |  |
| Figure 3 | a | 3367 | F | DTA<sup>fl/wt</sup>//Glast-CreERT<sup>wt/wt</sup> + Tamoxifen | 3 dpa |  |
|  |  | 314 | F | DTA<sup>fl/wt</sup>//Glast-CreERT<sup>tg/wt</sup> + Tamoxifen | 11 dpa |  |
|  | b,c | 3367 | F | DTA<sup>fl/wt</sup>//Glast-CreERT<sup>wt/wt</sup> + Tamoxifen | 3 dpa | n=3 (control) |
|  |  | 4902 | M | DTA<sup>wt/wt</sup>//Glast-CreERT<sup>tg/wt</sup> + Tamoxifen | 1 dpa |  |
|  |  | 4518 | F | DTA<sup>fl/wt</sup>//Glast-CreERT<sup>wt/wt</sup> + Tamoxifen | 3 dpa |  |
|  |  | 868 | M | DTA<sup>fl/wt</sup>//Glast-CreERT<sup>tg/wt</sup> + Tamoxifen | 6 hpa | n=3 (6 hpa) |
|  |  | 723 | F | DTA<sup>fl/wt</sup>//Glast-CreERT<sup>tg/wt</sup> + Tamoxifen | 6 hpa |  |
|  |  | 309 | F | DTA<sup>fl/wt</sup>//Glast-CreERT<sup>tg/wt</sup> + Tamoxifen | 6 hpa |  |
|  |  | 305 | M | DTA<sup>fl/wt</sup>//Glast-CreERT<sup>tg/wt</sup> + Tamoxifen | 11 dpa | n=3 (11 dpa) |
|  |  | 314 | F | DTA<sup>fl/wt</sup>//Glast-CreERT<sup>tg/wt</sup> + Tamoxifen | 11 dpa |  |
|  |  | 301 | F | DTA<sup>fl/wt</sup>//Glast-CreERT<sup>tg/wt</sup> + Tamoxifen | 11 dpa |  |
|  |  | 220 | F | DTA<sup>fl/wt</sup>//Glast-CreERT<sup>tg/wt</sup> + Tamoxifen | 28 dpa | n=3 (28 dpa) |
|  |  | 4836 | M | DTA<sup>fl/wt</sup>//Glast-CreERT<sup>tg/wt</sup> + Tamoxifen | 28 dpa |  |
|  |  | 271 | F | DTA<sup>fl/wt</sup>//Glast-CreERT<sup>tg/wt</sup> + Tamoxifen | 28 dpa |  |
|  | d | 3367 | F | DTA<sup>fl/wt</sup>//Glast-CreERT<sup>wt/wt</sup> + Tamoxifen | 3 dpa |  |
|  |  | 314 | F | DTA<sup>fl/wt</sup>//Glast-CreERT<sup>tg/wt</sup> + Tamoxifen | 11 dpa |  |
|  | e | 528 | F | DTA<sup>wt/wt</sup>//Glast-CreERT<sup>tg/wt</sup> + Tamoxifen | 6 hpa | n=3 (control) |
|  |  | 534 | M | DTA<sup>fl/wt</sup>//Glast-CreERT<sup>wt/wt</sup> + Tamoxifen | 11 dpa |  |
|  |  | 3367 | F | DTA<sup>fl/wt</sup>//Glast-CreERT<sup>wt/wt</sup> + Tamoxifen | 3 dpa |  |
|  |  | 309 | F | DTA<sup>fl/wt</sup>//Glast-CreERT<sup>tg/wt</sup> + Tamoxifen | 6 hpa | n=3 (6 hpa) |
|  |  | 868 | M | DTA<sup>fl/wt</sup>//Glast-CreERT<sup>tg/wt</sup> + Tamoxifen | 6 hpa |  |
|  |  | 723 | F | DTA<sup>fl/wt</sup>//Glast-CreERT<sup>tg/wt</sup> + Tamoxifen | 6 hpa |  |
|  |  | 305 | M | DTA<sup>fl/wt</sup>//Glast-CreERT<sup>tg/wt</sup> + Tamoxifen | 11 dpa | n=3 (11 dpa) |
|  |  | 310 | F | DTA<sup>fl/wt</sup>//Glast-CreERT<sup>tg/wt</sup> + Tamoxifen | 11 dpa |  |
|  |  | 314 | F | DTA<sup>fl/wt</sup>//Glast-CreERT<sup>tg/wt</sup> + Tamoxifen | 11 dpa |  |
|  |  | 217 | M | DTA<sup>fl/wt</sup>//Glast-CreERT<sup>tg/wt</sup> + Tamoxifen | 28 dpa | n=3 (28 dpa) |
|  |  | 4836 | M | DTA<sup>fl/wt</sup>//Glast-CreERT<sup>tg/wt</sup> + Tamoxifen | 28 dpa |  |
|  |  | 4631 | F | DTA<sup>fl/wt</sup>//Glast-CreERT<sup>tg/wt</sup> + Tamoxifen | 28 dpa |  |
| Figure 4 | a | 4224 | M | DTA<sup>fl/wt</sup>//Glast-CreERT<sup>wt/wt</sup> + Tamoxifen | 3 dpa |  |
|  |  | 294 | F | DTA<sup>fl/wt</sup>//Glast-CreERT<sup>tg/wt</sup> + Tamoxifen | 1 dpa |  |
|  |  | 314 | F | DTA<sup>fl/wt</sup>//Glast-CreERT<sup>tg/wt</sup> + Tamoxifen | 11 dpa |  |
|  |  | 4863 | M | DTA<sup>fl/wt</sup>//Glast-CreERT<sup>tg/wt</sup> + Tamoxifen | 28 dpa |  |
|  | b | 4862 | M | DTA<sup>fl/wt</sup>//Glast-CreERT<sup>wt/wt</sup> + Tamoxifen | 28 dpa |  |
|  |  | 293 | F | DTA<sup>fl/wt</sup>//Glast-CreERT<sup>tg/wt</sup> + Tamoxifen | 1 dpa |  |
|  |  | 310 | M | DTA<sup>fl/wt</sup>//Glast-CreERT<sup>tg/wt</sup> + Tamoxifen | 11 dpa |  |
|  |  | 4624 | M | DTA<sup>fl/wt</sup>//Glast-CreERT<sup>tg/wt</sup> + Tamoxifen | 28 dpa |  |
| Figure 5 | a | 4287 | F | DTA<sup>fl/wt</sup>//Glast-CreERT<sup>wt/wt</sup> + Tamoxifen | 1 dpa |  |
|  |  | 293 | M | DTA<sup>fl/wt</sup>//Glast-CreERT<sup>tg/wt</sup> + Tamoxifen | 1 dpa |  |
|  |  | 314 | F | DTA<sup>fl/wt</sup>//Glast-CreERT<sup>tg/wt</sup> + Tamoxifen | 11 dpa |  |
|  |  | 4631 | F | DTA<sup>fl/wt</sup>//Glast-CreERT<sup>tg/wt</sup> + Tamoxifen | 28 dpa |  |
|  | b-d,f,h | 4287 | F | DTA<sup>fl/wt</sup>//Glast-CreERT<sup>wt/wt</sup> + Tamoxifen | 1 dpa | n=3 (control) |
|  |  | 301 | F | DTA<sup>fl/wt</sup>//Glast-CreERT<sup>wt/wt</sup> + Tamoxifen | 11 dpa |  |
|  |  | 219 | F | DTA<sup>wt/wt</sup>//Glast-CreERT<sup>wt/wt</sup> + Tamoxifen | 28 dpa |  |
|  |  | 293 | M | DTA<sup>fl/wt</sup>//Glast-CreERT<sup>tg/wt</sup> + Tamoxifen | 1 dpa | n=3 (1 dpa) |
|  |  | 4221 | F | DTA<sup>fl/wt</sup>//Glast-CreERT<sup>tg/wt</sup> + Tamoxifen | 1 dpa |  |
|  |  | 4901 | M | DTA<sup>fl/wt</sup>//Glast-CreERT<sup>tg/wt</sup> + Tamoxifen | 1 dpa |  |
|  |  | 271 | F | DTA<sup>fl/wt</sup>//Glast-CreERT<sup>tg/wt</sup> + Tamoxifen | 11 dpa | n=3 (11 dpa) |
|  |  | 310 | M | DTA<sup>fl/wt</sup>//Glast-CreERT<sup>tg/wt</sup> + Tamoxifen | 11 dpa |  |
|  |  | 314 | F | DTA<sup>fl/wt</sup>//Glast-CreERT<sup>tg/wt</sup> + Tamoxifen | 11 dpa |  |
|  |  | 291 | F | DTA<sup>fl/wt</sup>//Glast-CreERT<sup>tg/wt</sup> + Tamoxifen | 28 dpa | n= 3 (28 dpa) |
|  |  | 4631 | F | DTA<sup>fl/wt</sup>//Glast-CreERT<sup>tg/wt</sup> + Tamoxifen | 28 dpa |  |
|  |  | 4865 | M | DTA<sup>fl/wt</sup>//Glast-CreERT<sup>tg/wt</sup> + Tamoxifen | 28 dpa |  |
| Supp. Fig. 1 | a-d | 4864 | M | DTA<sup>fl/wt</sup>//Glast-CreERT<sup>wt/wt</sup> + Tamoxifen | 28 dpa |  |
|  |  | 314 | F | DTA<sup>fl/wt</sup>//Glast-CreERT<sup>tg/wt</sup> + Tamoxifen | 11 dpa |  |
| Supp. Fig. 2 | a | 381 | M | DTA<sup>fl/wt</sup>//Glast-CreERT<sup>tg/wt</sup>, Naïve | N/A |  |
|  |  | 382 | F | DTA<sup>fl/wt</sup>//Glast-CreERT<sup>tg/wt</sup>, Naïve | N/A |  |
|  |  | 4525 | M | DTA<sup>fl/wt</sup>//Glast-CreERT<sup>tg/wt</sup> + Oil | 3 dpa |  |
|  | b,c | 4862 | M | DTA<sup>fl/wt</sup>//Glast-CreERT<sup>wt/wt</sup> + Tamoxifen | 28 dpa |  |
|  |  | 4864 | M | DTA<sup>fl/wt</sup>//Glast-CreERT<sup>wt/wt</sup> + Tamoxifen | 28 dpa |  |
|  |  | 305 | M | DTA<sup>fl/wt</sup>//Glast-CreERT<sup>tg/wt</sup> + Tamoxifen | 11 dpa |  |
|  | d | 4862 | M | DTA<sup>fl/wt</sup>//Glast-CreERT<sup>wt/wt</sup> + Tamoxifen | 28 dpa |  |
|  |  | 305 | M | DTA<sup>fl/wt</sup>//Glast-CreERT<sup>tg/wt</sup> + Tamoxifen | 11 dpa |  |
| Supp. Fig. 3 | a,b | 292 | F | DTA<sup>fl/wt</sup>//Glast-CreERT<sup>wt/wt</sup> + Tamoxifen | 1 dpa |  |
|  |  | 892 | M | DTA<sup>fl/wt</sup>//Glast-CreERT<sup>tg/wt</sup> + Tamoxifen | 2 hpa |  |
|  |  | 4624 | M | DTA<sup>fl/wt</sup>//Glast-CreERT<sup>tg/wt</sup> + Tamoxifen | 28 dpa |  |
| Supp. Fig. 4 | a | 4902 | M | DTA<sup>wt/wt</sup>//Glast-CreERT<sup>tg/wt</sup> + Tamoxifen | 1 dpa |  |
|  |  | 4519 | M | DTA<sup>fl/wt</sup>//Glast-CreERT<sup>tg/wt</sup> + Tamoxifen | 3 dpa |  |
| Supp. Fig. 5 |  | 4225 | M | DTA<sup>wt/wt</sup>//Glast-CreERT<sup>tg/wt</sup> + Tamoxifen | 1 dpa |  |
|  |  | 4223 | F | DTA<sup>fl/wt</sup>//Glast-CreERT<sup>tg/wt</sup> + Tamoxifen | 1 dpa |  |
|  |  | 4519 | M | DTA<sup>fl/wt</sup>//Glast-CreERT<sup>tg/wt</sup> + Tamoxifen | 3 dpa |  |
|  |  | 314 | F | DTA<sup>fl/wt</sup>//Glast-CreERT<sup>tg/wt</sup> + Tamoxifen | 11 dpa |  |
|  |  | 4863 | M | DTA<sup>fl/wt</sup>//Glast-CreERT<sup>tg/wt</sup> + Tamoxifen | 28 dpa |  |
| Supp. Fig. 6 | a | 2794 | M | DTA<sup>wt/wt</sup>//Glast-CreERT<sup>tg/wt</sup> + Tamoxifen | 3 dpa |  |
|  | b | 537 | M | DTA<sup>fl/wt</sup>//Glast-CreERT<sup>wt/wt</sup> + Tamoxifen | 5 dpa |  |
|  |  | 4519 | M | DTA<sup>fl/wt</sup>//Glast-CreERT<sup>tg/wt</sup> + Tamoxifen | 3 dpa |  |
|  |  | 305 | M | DTA<sup>fl/wt</sup>//Glast-CreERT<sup>tg/wt</sup> + Tamoxifen | 11 dpa |  |
|  |  | 4631 | F | DTA<sup>fl/wt</sup>//Glast-CreERT<sup>tg/wt</sup> + Tamoxifen | 28 dpa |  |
| Supp.<br>Fig. 7 | c | 2794 | M | DTA<sup>wt/wt</sup>//Glast-CreERT<sup>tg/wt</sup> + Tamoxifen | 3 dpa |  |
|  |  | 4287 | F | DTA<sup>fl/wt</sup>//Glast-CreERT<sup>tg/wt</sup> + Tamoxifen | 1 dpa |  |
|  |  | 2799 | M | DTA<sup>fl/wt</sup>//Glast-CreERT<sup>tg/wt</sup> + Tamoxifen | 3 dpa |  |
|  |  | 535 | F | DTA<sup>fl/wt</sup>//Glast-CreERT<sup>tg/wt</sup> + Tamoxifen | 5 dpa |  |
| In-text<br>(Results<br>Section<br>3.7) | **Ki67** | 897 | M | DTA<sup>fl/wt</sup>//Glast-CreERT<sup>wt/wt</sup> + Tamoxifen | 2 hpa | n=10 (control) |
|  |  | 0043 | F | DTA<sup>fl/wt</sup>//Glast-CreERT<sup>wt/wt</sup> + Tamoxifen | 1 dpa |  |
|  |  | 4224 | M | DTA<sup>fl/wt</sup>//Glast-CreERT<sup>wt/wt</sup> + Tamoxifen | 3 dpa |  |
|  |  | 4225 | M | DTA<sup>fl/fl</sup>//Glast-CreERT<sup>wt/wt</sup> + Tamoxifen | 1 dpa |  |
|  |  | 4233 | M | DTA<sup>fl/wt</sup>//Glast-CreERT<sup>wt/wt</sup> + Tamoxifen | 1 dpa |  |
|  |  | 4287 | M | DTA<sup>fl/wt</sup>//Glast-CreERT<sup>wt/wt</sup> + Tamoxifen | 1 dpa |  |
|  |  | 2794 | M | DTA<sup>wt/wt</sup>//Glast-CreERT<sup>tg/wt</sup> + Tamoxifen | 3 dpa |  |
|  |  | 3359 | M | DTA<sup>wt/wt</sup>//Glast-CreERT<sup>tg/wt</sup> + Tamoxifen | 3 dpa |  |
|  |  | 3360 | F | DTA<sup>wt/wt</sup>//Glast-CreERT<sup>tg/wt</sup> + Tamoxifen | 3 dpa |  |
|  |  | 4864 | M | DTA<sup>fl/wt</sup>//Glast-CreERT<sup>wt/wt</sup> + Tamoxifen | 28 dpa |  |
|  |  | 893 | F | DTA<sup>fl/wt</sup>//Glast-CreERT<sup>tg/wt</sup> + Tamoxifen | 2 hpa | n=3 (2 hpa) |
|  |  | 895 | M | DTA<sup>fl/wt</sup>//Glast-CreERT<sup>tg/wt</sup> + Tamoxifen | 2 hpa |  |
|  |  | 892 | M | DTA<sup>fl/wt</sup>//Glast-CreERT<sup>tg/wt</sup> + Tamoxifen | 2 hpa |  |
|  |  | 309 | F | DTA<sup>fl/wt</sup>//Glast-CreERT<sup>tg/wt</sup> + Tamoxifen | 6 hpa | n=3 (6 hpa) |
|  |  | 868 | M | DTA<sup>fl/wt</sup>//Glast-CreERT<sup>tg/wt</sup> + Tamoxifen | 6 hpa |  |
|  |  | 723 | F | DTA<sup>fl/wt</sup>//Glast-CreERT<sup>tg/wt</sup> + Tamoxifen | 6 hpa |  |
|  |  | 0046 | M | DTA<sup>fl/wt</sup>//Glast-CreERT<sup>tg/wt</sup> + Tamoxifen | 1 dpa | n=3 (1 dpa) |
|  |  | 3497 | F | DTA<sup>fl/wt</sup>//Glast-CreERT<sup>tg/wt</sup> + Tamoxifen | 1 dpa |  |
|  |  | 4221 | F | DTA<sup>fl/wt</sup>//Glast-CreERT<sup>tg/wt</sup> + Tamoxifen | 1 dpa |  |
|  |  | 4223 | F | DTA<sup>fl/wt</sup>//Glast-CreERT<sup>tg/wt</sup> + Tamoxifen | 1 dpa |  |
|  |  | 2799 | M | DTA<sup>fl/wt</sup>//Glast-CreERT<sup>tg/wt</sup> + Tamoxifen | 3 dpa | n=5 (3 dpa) |
|  |  | 3352 | M | DTA<sup>fl/wt</sup>//Glast-CreERT<sup>tg/wt</sup> + Tamoxifen | 3 dpa |  |
|  |  | 3357 | F | DTA<sup>fl/wt</sup>//Glast-CreERT<sup>tg/wt</sup> + Tamoxifen | 3 dpa |  |
|  |  | 3365 | F | DTA<sup>fl/wt</sup>//Glast-CreERT<sup>tg/wt</sup> + Tamoxifen | 3 dpa |  |
|  |  | 4519 | M | DTA<sup>fl/wt</sup>//Glast-CreERT<sup>tg/wt</sup> + Tamoxifen | 3 dpa |  |
|  |  | 535 | F | DTA<sup>fl/wt</sup>//Glast-CreERT<sup>tg/wt</sup> + Tamoxifen | 5 dpa | n=3 (5 dpa) |
|  |  | 536 | M | DTA<sup>fl/wt</sup>//Glast-CreERT<sup>tg/wt</sup> + Tamoxifen | 5 dpa |  |
|  |  | 538 | M | DTA<sup>fl/wt</sup>//Glast-CreERT<sup>tg/wt</sup> + Tamoxifen | 5 dpa |  |
|  |  | 305 | M | DTA<sup>fl/wt</sup>//Glast-CreERT<sup>tg/wt</sup> + Tamoxifen | 11 dpa | n=3 (11 dpa) |
|  |  | 310 | M | DTA<sup>fl/wt</sup>//Glast-CreERT<sup>tg/wt</sup> + Tamoxifen | 11 dpa |  |
|  |  | 314 | F | DTA<sup>fl/wt</sup>//Glast-CreERT<sup>tg/wt</sup> + Tamoxifen | 11 dpa |  |
|  |  | 4624 | M | DTA<sup>fl/wt</sup>//Glast-CreERT<sup>tg/wt</sup> + Tamoxifen | 28 dpa | n=4 (28 dpa) |
|  |  | 4631 | F | DTA<sup>fl/wt</sup>//Glast-CreERT<sup>tg/wt</sup> + Tamoxifen | 28 dpa |  |
|  |  | 4836 | M | DTA<sup>fl/wt</sup>//Glast-CreERT<sup>tg/wt</sup> + Tamoxifen | 28 dpa |  |
|  |  | 4865 | M | DTA<sup>fl/wt</sup>//Glast-CreERT<sup>tg/wt</sup> + Tamoxifen | 28 dpa |  |
|  | **BrdU** | 897 | M | DTA<sup>fl/wt</sup>//Glast-CreERT<sup>wt/wt</sup> + Tamoxifen | 2 hpa | n=9 (control) |
|  |  | 319 | F | DTA<sup>wt/wt</sup>//Glast-CreERT<sup>tg/wt</sup> + Tamoxifen | 6 hpa |  |
|  |  | 4224 | M | DTA<sup>fl/wt</sup>//Glast-CreERT<sup>wt/wt</sup> + Tamoxifen | 3 dpa |  |
|  |  | 4225 | M | DTA<sup>fl/fl</sup>//Glast-CreERT<sup>wt/wt</sup> + Tamoxifen | 1 dpa |  |
|  |  | 4287 | F | DTA<sup>fl/wt</sup>//Glast-CreERT<sup>wt/wt</sup> + Tamoxifen | 1 dpa |  |
|  |  | 2794 | M | DTA<sup>wt/wt</sup>//Glast-CreERT<sup>tg/wt</sup> + Tamoxifen | 3 dpa |  |
|  |  | 3359 | M | DTA<sup>wt/wt</sup>//Glast-CreERT<sup>tg/wt</sup> + Tamoxifen | 3 dpa |  |
|  |  | 3360 | F | DTA<sup>wt/wt</sup>//Glast-CreERT<sup>tg/wt</sup> + Tamoxifen | 3 dpa |  |
|  |  | 537 | M | DTA<sup>fl/wt</sup>//Glast-CreERT<sup>wt/wt</sup> + Tamoxifen | 5 dpa |  |
|  |  | 893 | F | DTA<sup>fl/wt</sup>//Glast-CreERT<sup>tg/wt</sup> + Tamoxifen | 2 hpa | n=3 (2 hpa) |
|  |  | 895 | M | DTA<sup>fl/wt</sup>//Glast-CreERT<sup>tg/wt</sup> + Tamoxifen | 2 hpa |  |
|  |  | 892 | M | DTA<sup>fl/wt</sup>//Glast-CreERT<sup>tg/wt</sup> + Tamoxifen | 2 hpa |  |
|  |  | 309 | F | DTA<sup>fl/wt</sup>//Glast-CreERT<sup>tg/wt</sup> + Tamoxifen | 6 hpa | n=3 (6 hpa) |
|  |  | 868 | M | DTA<sup>fl/wt</sup>//Glast-CreERT<sup>tg/wt</sup> + Tamoxifen | 6 hpa |  |
|  |  | 723 | F | DTA<sup>fl/wt</sup>//Glast-CreERT<sup>tg/wt</sup> + Tamoxifen | 6 hpa |  |
|  |  | 0046 | M | DTA<sup>fl/wt</sup>//Glast-CreERT<sup>tg/wt</sup> + Tamoxifen | 1 dpa | n=3 (1 dpa) |
|  |  | 3497 | F | DTA<sup>fl/wt</sup>//Glast-CreERT<sup>tg/wt</sup> + Tamoxifen | 1 dpa |  |
|  |  | 4221 | F | DTA<sup>fl/wt</sup>//Glast-CreERT<sup>tg/wt</sup> + Tamoxifen | 1 dpa |  |
|  |  | 4223 | F | DTA<sup>fl/wt</sup>//Glast-CreERT<sup>tg/wt</sup> + Tamoxifen | 1 dpa |  |
|  |  | 2799 | M | DTA<sup>fl/wt</sup>//Glast-CreERT<sup>tg/wt</sup> + Tamoxifen | 3 dpa | n=5 (3 dpa) |
|  |  | 3352 | M | DTA<sup>fl/wt</sup>//Glast-CreERT<sup>tg/wt</sup> + Tamoxifen | 3 dpa |  |
|  |  | 3357 | F | DTA<sup>fl/wt</sup>//Glast-CreERT<sup>tg/wt</sup> + Tamoxifen | 3 dpa |  |
|  |  | 3365 | F | DTA<sup>fl/wt</sup>//Glast-CreERT<sup>tg/wt</sup> + Tamoxifen | 3 dpa |  |
|  |  | 4519 | M | DTA<sup>fl/wt</sup>//Glast-CreERT<sup>tg/wt</sup> + Tamoxifen | 3 dpa |  |
|  |  | 535 | F | DTA<sup>fl/wt</sup>//Glast-CreERT<sup>tg/wt</sup> + Tamoxifen | 5 dpa | n=3 (5 dpa) |
|  |  | 536 | F | DTA<sup>fl/wt</sup>//Glast-CreERT<sup>tg/wt</sup> + Tamoxifen | 5 dpa |  |
|  |  | 538 | M | DTA<sup>fl/wt</sup>//Glast-CreERT<sup>tg/wt</sup> + Tamoxifen | 5 dpa |  |

Diphtheria Toxin fragment A (DTA) interferes with protein synthesis, inducing apoptotic cell death in those cells that express it (Ivanova, Signore et al. 2005). DTA^fl/wt^ (fl: floxed allele, encoding loxP sites; wt: wildtype allele) mice express DTA under the Rosa26 promoter behind a stop cassette flanked by loxP sites, which interferes with DTA expression until the stop cassette is removed by Cre recombinase. Cre recombinase is fused with an estrogen receptor and expressed behind the Glast promoter and as part of a complex in Glast-CreERT^tg/wt^ mice (tg: allele carries transgene; wt: wildtype allele). This restricts expression to astrocytes and enables timed relocalization of the CreERT protein complex to the nucleus for excision of the stop cassette only after Tamoxifen (TX) administration (**Fig. 1a**). Glast-CreERT is expressed and causes Cre-mediated recombination in a limited number of astrocytes in the forebrain of adult mice (Mori et al., 2006; Srinivasan et al., 2016).

**Figure 1.**
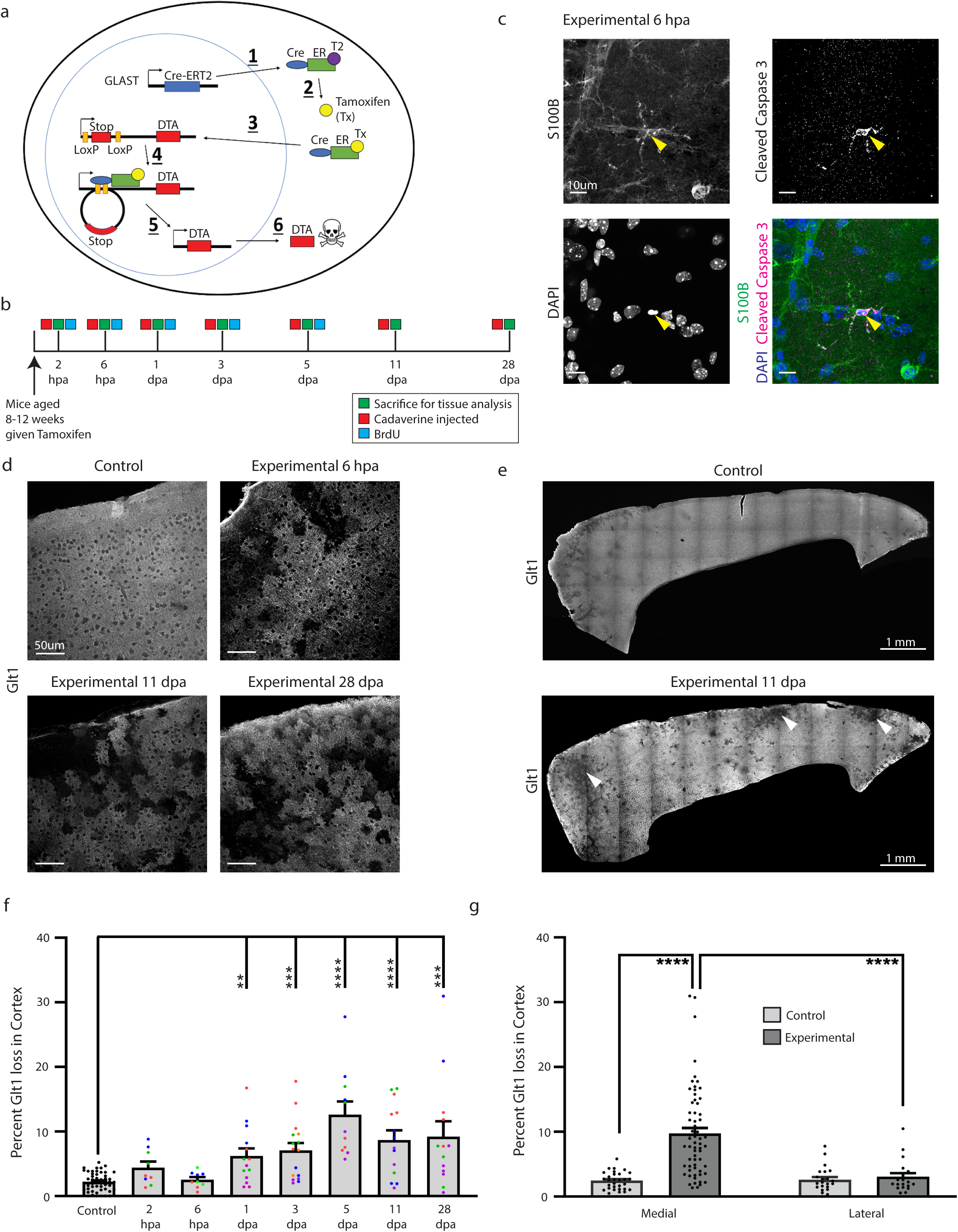
Astrocyte ablation occurred within hours after Tamoxifen administration. **a.** Astrocytes were ablated in mice using a cre-inducible system expressing diptheria toxin fragment fragment a (DTA) after a stop cassette flanked by loxP sites: 1) Cre recombinase coupled to an estrogen receptor was expressed behind the astrocyte glutamate transporter GLAST. 2) Tamoxifen was administered and binded to this estrogen receptor, enabling 3) translocation of cre into the nucleus where 4) recombination at LoxP sites excised the stop sequence in front of DTA and 5) enabled DTA transcription and translation. 6) DTA halted protein translation and led to apoptotic cell death. **b.** Mice were sacrificed at multiple timepoints between 2 hours and 28 days after tamoxifen administration to determine the timeline and extent of astrocyte ablation. Some animals were also injected with Cadaverine and/or BrdU. **c.** Some astrocytes in DTA^fl/wt^//Glast-CreERT^tg/wt^ mice colocalized with the apoptotic marker cleaved caspase 3 shortly after tamoxifen administration (arrowhead). **d.** Loss of the astrocyte marker Glt1 indicating cell ablation occurred only in DTA^fl/wt^//Glast-CreERT^tg/wt^ mice several hours after tamoxifen administration and continued up to 28 days later, while no Glt1 loss was detected at any timepoint in DTA^fl/wt^//Glast-CreERT^wt/wt^ or DTA^wt/wt^//Glast-CreERT^tg/wt^ mice given tamoxifen. **e.** Glt1 loss indicative of astrocyte ablation occurred across the cortex in DTA^fl/wt^//Glast-CreERT^tg/wt^ mice after tamoxifen administration (arrowheads). DTA^fl/wt^//Glast-CreERT^wt/wt^ or DTA^wt/wt^//Glast-CreERT^tg/wt^ mice showed few “Glt1-low regions” along the midline or along blood vessels. **f.** The percentage of Glt1-negative areas were quantified in the cortical gray matter per slice across all timepoints. Slices from the same animal were plotted in the same color. **g.** Glt1 loss percentage for each slice were plotted in either medial or lateral groups based on location in *The Allen Brain Atlas*.

Experimental mice had the following genotype: DTA^fl/wt^//Glast-CreERT^tg/w^, and received a single dose of Tamoxifen. Controls were as follows: 1. Some DTA^fl/wt^//Glast-CreERT^tg/wt^ mice received carrier solution only, which lacked tamoxifen. 2. Mice lacking either the DTA or the Glast-CreERT allele (DTA^fl/wt^//Glast-CreERT^wt/wt^ or DTA^wt/wt^//Glast-CreERT^tg/wt^) received Tamoxifen. 3. Naive DTA^fl/wt^//Glast-CreERT^tg/wt^ mice received no treatment. No differences between control groups were detected and controls were hence pooled for analysis.

42 experimental (DTA^fl/wt^//Glast-CreERT^tg/wt^) and 60 mice (DTA^fl/wt^//Glast-CreER^wt/wt^ or DTA^wt/wt^//Glast-CreERT^tg/wt^) received a single dose of Tamoxifen at 8-12 weeks of age. Mice were injected with a lethal dose of Ketamine (100 mg/kg) /Xylazine (10 mg/kg) followed by transcardial perfusion for subsequent histology at 2 and 6 hours post ablation (hpa) as well as 1, 3, 5, 11 and 28 days post ablation (dpa) (**Fig. 1b**). Some mice were injected retro-orbitally with Cadaverine to assess blood-brain barrier integrity.

### 2.3 Tamoxifen administration

Tamoxifen (Sigma, cat #T6548) was dissolved at 40 mg/mL in a mixture of 10% 200-proof ethanol and 90% Corn oil (Sigma, cat #C2867) for 2 h while shaking at 37°C. The tamoxifen solution was administered via oral gavage at 330 mg/kg once.

### 2.4 Cadaverine administration

To assess blood-brain barrier function Alexa Fluor-555 Cadaverine (950 Da,1 mg; Invitrogen, Catalog #A30677) was injected into the retro-orbital sinus of each mouse. Cadaverine (1 mg) was dissolved in 300 uL of sterile saline. Each mouse was injected with a volume of 100 uL (0.33 mg). Mice were perfused transcardially with Phospho-Buffered Saline (PBS) followed by 4% paraformaldehyde (PFA) 30 minutes after Cadaverine injection.

### 2.5 BrdU administration

For the assessment of astrocyte proliferation, intraperitoneal injections of BrdU/saline (50μg/g body weight) were administered to experimental and control mice twice daily for 5 days, starting on the day of Tx administration. Mice were sacrificed at 2 hpa, 6 hpa, 1 dpa, 3 dpa, or 5 dpa.

### 2.6 Histology

Brains were collected after transcardial perfusion and post-fixed in 4% PFA overnight. Sagittal slices were cut at 50 μm thickness using a vibratome (Campden 5100mz). Immunohistochemistry used the primary antibodies (listed in **Table 2**) in PBS with 10% goat serum 0.5% Triton X-100 at 4°C overnight. Slices were washed in PBS and incubated in secondary antibody (listed in **Table 2**) solution of PBS with 10% goat serum and 0.5% Triton X-100 for 1-2 h at room temperature. 4,6-diamidino-2-phenylindole (DAPI) was included in the secondary antibody solution as needed. Slices were washed in PBS three times for ten minutes each and then mounted onto glass microscope slides with Aqua Poly/Mount (Polysciences, catalog #18606).

**Table 2.** Antibodies used in experiments.

| Primary Antibodies |  |  |  |  |  |  |
| --- | --- | --- | --- | --- | --- | --- |
| Name | Manufacturer | Catalog # | RRID | Species Raised in | Monoclonal/<br>Polyclonal | Concentration |
| S100β | Sigma Aldrich | S2532 | AB_477499 | Mouse | Monoclonal | 1 : 1000 |
| Glt1 | Millipore | AB1783 | AB_90949 | Guinea pig | Polyclonal | 1 : 1000 |
| GFAP | Millipore | AB5541 | AB_177521 | Chicken | Polyclonal | 1 : 1000 |
| Ki67 | Thermo Fisher | RM-9106-S1 | AB_149792 | Rabbit | Monoclonal | 1 : 1000 |
| BrdU | Abcam | ab6326 | AB_305426 | Rat | Monoclonal | 1 : 500 |
| Cleaved<br>Caspase 3 | Cell Signaling<br>Technology | 9661 | AB_2341188 | Rabbit | Polyclonal | 1 : 1000 |
| Sox-9 | Millipore | AB5535 | AB_2239761 | Rabbit | Polyclonal | 1 : 1000 |
| Fibrinogen | Agilent | A008002 | AB_578481 | Rabbit | Polyclonal | 1 : 500 |
| ZO-1 | Abcam | ab96587 | AB-18.0006 | Rabbit | Polyclonal | 1 : 100 |
| pSTAT3 | Cell Signaling<br>Technology | #9145S | AB_2491009 | Rabbit | Monoclonal | 1:100 |
| Iba1 | Wako | #09-19741 | AB_839504 | Rabbit | Polyclonal | 1:1000 |

| Secondary Antibodies |  |  |  |  |  |  |
| --- | --- | --- | --- | --- | --- | --- |
| Name | Manufacturer | Catalog # | RRID | Species Raised in | Monoclonal/ Polyclonal | Concentration |
| Chicken Alexa-488 | Jackson Immuno Research | 703-546-155 | AB_2340376 | Donkey | Polyclonal | 1 : 1000 |
| Rabbit Alexa-488 | Jackson Immuno Research | 111-546-144 | AB_2338057 | Donkey | Polyclonal | 1 : 1000 |
| Mouse Alexa-488 | Jackson Immuno Research | 115-546-003 | AB_2338859 | Goat | Polyclonal | 1 : 1000 |
| Rat Alexa-488 | Jackson Immuno Research | 112-546-003 | AB_2338364 | Goat | Polyclonal | 1 : 1000 |
| Guinea pig Alexa-647 | Jackson Immuno Research | 106-606-003 | AB_2337449 | Goat | Polyclonal | 1 : 1000 |

| Dyes |  |  |  |  |  |  |
| --- | --- | --- | --- | --- | --- | --- |
| Name | Manufacturer | Catalog # | RRID | Species Raised in | Monoclonal/ Polyclonal | Concentration |
| DAPI | ThermoFisher | D1306 | AB_2629482 | N /A | N /A | 1 : 1000 |
| Alexa-555 Cadaverine | ThermoFisher | A30677 | N/A | N/A | N/A | 0.33mg/mouse |

For immunohistochemistry against BrdU, antigen retrieval was performed using 1x citrate buffer (pH 6.0,Thermo Fisher, Catalog #005000) at 95°C for 20 minutes followed by incubation in primary antibody solution overnight at 4°C. This was followed by one 10 minute wash in 0.5% Triton X/ PBS and two 10 minute washes in PBS. Slices were mounted on glass slides, covered with Aqua Poly/Mount and a glass coverslip. Slices stained for phosphorylated Stat-3 underwent antigen retrieval in 10 mM Tris-HCl, 1 mM EDTA at pH 9.0 at 90°C for 20 minutes. Slices were washed in PBS for 1 hour before incubating in primary antibody solution overnight at 4°C.

For tight junction (ZO-1) staining, slices were subjected to antigen retrieval using 100 mg pepsin in 10mM hydrochloric acid (HCl) for 20 minutes at 37°C. Next, slices were washed twice in PBST (PBS with 150 uL L^-1^ Tween-20) and then incubated in 3% H_2_O_2_ for 10 minutes. Finally, slices were washed 3 times in PBS for ten minutes each and then incubated in primary antibody solution for a minimum of 48 hours at 4°C. Images of mice were taken using a Nikon A1R confocal microscope with Nikon 4x, 10x or 20x air objectives or Nikon Apo 40x/1.30 and 60x/1.40 oil immersion objectives.

### 2.7 Data Analysis

#### 2.7.1 Quantification of astrocyte loss

Astrocyte loss was assessed in the cortical gray matter of experimental and control mice using immunohistochemistry against the astrocytic glutamate transporter Glt1, which is expressed in the fine processes in all astrocytes. Large image scans were taken of the cortex and/or hippocampus using a 20x objective and 2x line averaging at a single step position. Images were stitched together using optimal path stitching with 5% overlap. Line averaging was crucial to reduce grid lines where images overlap. Three to five sagittal slices per animal and 3 mice per group were imaged for quantification. Areas that lacked Glt1 labeling were identified after binarization of the image using set thresholding parameters in ImageJ. The binarized image was then processed using the “fill holes” feature to consolidate Glt1 loss areas into discrete shapes. Finally, Glt1 loss regions were found by setting consistent size and circularity parameters that excluded spaces covered by blood vessels and neurons from being recognized as “Glt1-free”. (**Suppl. Fig. 1**). Glt1 loss was calculated as a percentage of the total cortex area in each slice and was plotted with each data point representing one slice. Data points of slices from the same animal were plotted using the same color, different colors represent data points from different mice.

#### 2.7.2 Quantification of Cadaverine leakage

Cadaverine is coupled to the fluorescent dye Alexa Fluor 550 (A550) and is excluded from the brain parenchyma by the functional BBB. BBB leakiness permits the dye to cross into the brain, where neurons take it up. Large image scans of the cortex were taken using a 10x objective and images were stitched together using optimal path stitching with 5% overlap. Five slices were imaged per animal. Cadaverine^+^ regions were included in quantification upon meeting the following criteria: 1) Cadaverine cells were brighter than the background, 2) were clustered in close proximity to each other, and 3) each drawn ROI consisted of at least 10 Cadaverine^+^ cells. Cadaverine leakage area was calculated as a percentage of the total cortex area in each slice and was plotted by animal.

#### 2.7.3 Quantification of astrocyte process length and volume

To determine if astrocytes adjacent to ablated neighbors respond by changing their morphology, we employed the Simple Neurite Tracer (Longair, Baker, & Armstrong, 2011) in Fiji/ImageJ in order to create a 3D representation of an astrocyte and its processes. Total process length and volume of GFAP^+^ astrocyte processes were measured and compared, and Sholl analysis was employed to determine astrocyte process complexity. Early (1 dpa), middle (11 dpa) and late (28 dpa) timepoints were examined in three mice per timepoint. Sample size for each group was n=15 astrocytes. Five astrocytes were located across at least three different slices for each animal, and these three slices were chosen to represent different areas of the cortex. GFAP expression was used to analyze astrocytes and their major process length and thickness, and Glt-1 was used to confirm loss areas in the experimental mice.

#### 2.7.4 Polarity quantification

To investigate the polarity of the astrocytes adjacent to areas of loss, we first identified the relevant area of loss and drew a line between the two ends of this loss (Line 1) **(Fig. 5e)**. From the center of Line 1, Line 2 (the bisector) was drawn through the center of the soma of the astrocyte and extended to the end of the longest process in that direction. Next, Line 3 was drawn perpendicular to Line 2, and it also went through the center of the soma of the astrocyte and extended to the end of the longest process in that direction. Line 2 and Line 3 served to divide the astrocyte into quadrants, with the more proximal quadrants being the ones directly adjacent to the relevant loss area while the distal quadrants were the ones further removed from the loss.

**Figure 5.**
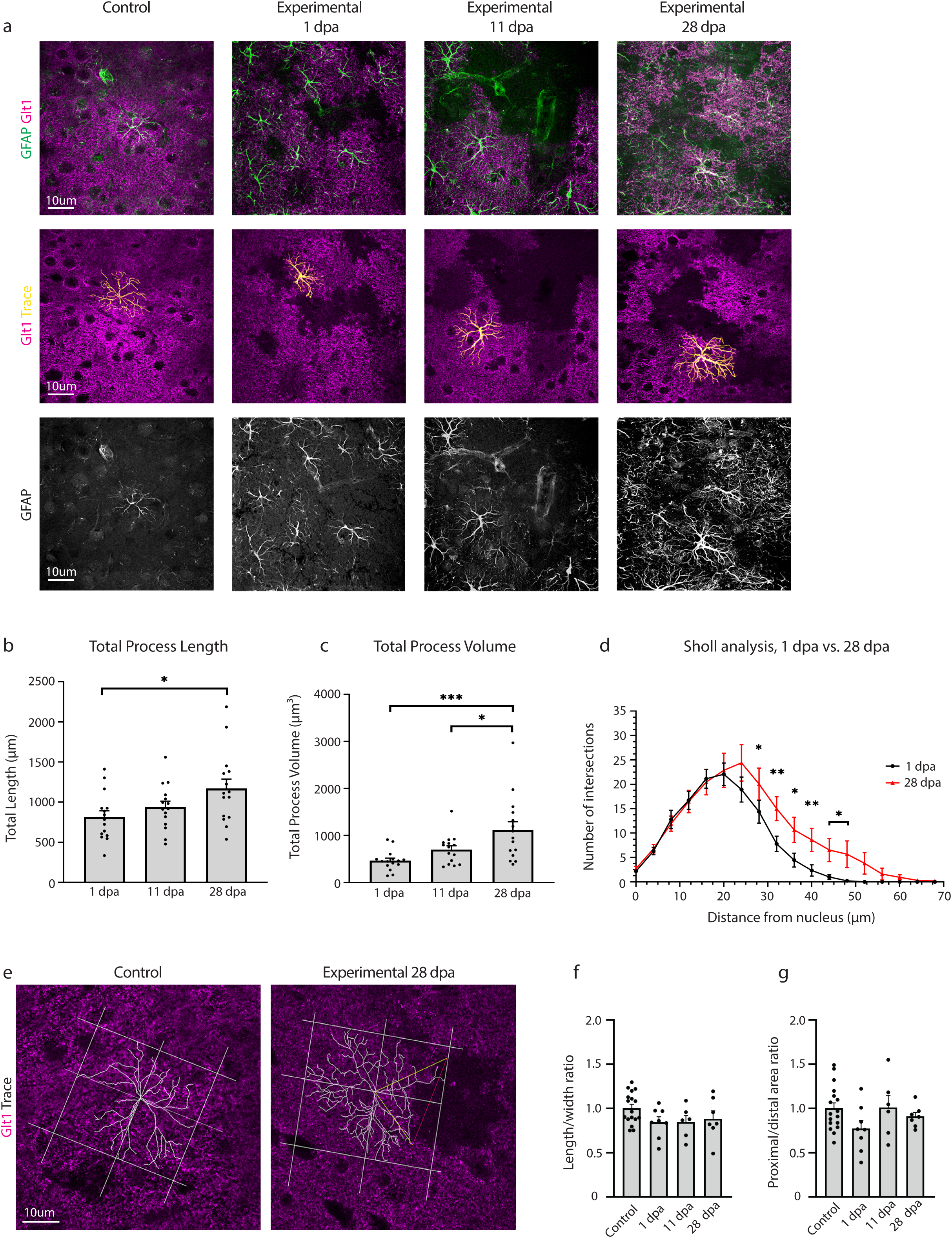
Astrocyte ablation induced GFAP upregulation in processes and mild morphological changes in neighboring astrocytes. **a.** Astrocytes adjacent to regions of astrocyte ablation indicated by lack of Glt1 showed increased expression of GFAP as early as 1 dpa and continued through 28 dpa. GFAP labeling was used to trace loss-adjacent astrocyte processes in ImageJ. **b.** Total length of all GFAP^+^ processes was quantified for loss-adjacent astrocytes at 1, 11, and 28 days post administration of tamoxifen. **c.** Total volume of all GFAP+ processes was quantified for loss-adjacent astrocytes at 1, 11, and 28 days post administration of tamoxifen. **d.** Sholl analysis compared arborization complexity of loss-adjacent astrocytes at 1 and 28 dpa. *p ≤ 0.05, **p≤ 0.01, ***p< 0.001, ****p ≤ 0.0001. **e.** Changes in astrocyte polarity were tested by drawing quadrants using astrocyte ablation regions as a reference when present. The two quadrants closest to ablation regions were called proximal, while the two quadrants furthest away were called distal. **f.** Length to width ratios of drawn quadrants were calculated and compared at 1, 11, and 28 dpa. **g.** The ratio of proximal quadrant areas to distal quadrant areas were calculated and compared among early, middle, and late timepoints.

Additional lines were added to enclose the quadrants, which were all centered at the astrocyte soma. The area of the two proximal quadrants and the area of the two distal quadrants was determined. The ratio of the distal area to the proximal area was then calculated, in order to provide us with information about the polarity of the astrocyte in relation to the relevant area of loss. The ratio of width (Line 3) to length (Line 2) was also calculated, in order to provide further polarity information.

For sham astrocytes, without a relevant area of loss to draw Line 1, a random number generator between 1 and 360 was used to determine the angle at which Line 1 would be drawn. From there, the same protocol from above was followed in order to divide the sham astrocytes into four quadrants. The ratio of the proximal area to the distal area and the ratio of width to length was calculated for these astrocytes as well.

#### 2.7.5 Quantification of astrocyte proliferation

Astrocyte proliferation was quantified using immunohistochemistry against the cell cycle protein Ki67 across all timepoints or BrdU labeling at 2 hpa, 6 hpa, 1 dpa, 3 dpa, and 5 dpa. To quantify the number of Ki67^+^ astrocytes, 1 confocal image was taken in 3-5 different slices per animal, 3 mice per group. To quantify the number of BrdU^+^ astrocytes, 1 confocal image was taken in 3-5 different slices per animal, 3 mice per group. For experimental mice, images were taken in areas adjacent to Glt1 loss. For control mice, one image each was taken in the frontal, medial, and lateral regions of cortex. Regions surrounding large penetrating arteries were avoided in both groups because Glt1 expression levels are sometimes reduced in directly adjacent astrocytes, even in controls. Confocal images had a z-step size of 1μm. Stacks spanned the entire slice and were quantified step-by-step using the cell counter tool in ImageJ. Cells that co-labeled for S100β, DAPI and Ki67 or S100β, DAPI and BrdU were considered proliferating astrocytes.

#### 2.7.6 Quantification of ZO-1 expression

Expression of the tight junction protein ZO-1 was quantified in ImageJ. ZO-1 images were binarized based on intensity and size and resulting masks were overlaid onto corresponding images of the marker CD31 that labels endothelial cells. A line was drawn along the CD31^+^ blood vessels, and the generated plot profile represented ZO-1-positive pixels as “1”, and pixels lacking ZO-1 as “0”. The plot profile data was used to calculate the percent of pixels in the drawn line that lacked ZO-1. Using the known pixel dimensions, this measurement could be converted into length of ZO-1 coverage in micrometers and was reported per animal as percent of vessel lacking ZO-1. 3-5 confocal images were taken in 3-5 different slices, 3 mice per group. The experimental mice were examined along early (6 hpa), middle (11 dpa) and late (28 dpa) timepoints. Images for experimental mice were taken in regions lacking Glt1. For the same ZO-1 images, fluorescence intensity was also measured and was reported per animal.

#### 2.7.7 Quantification of GLUT1 expression

We measured fluorescence intensity of the protein Glucose Transporter 1 (GLUT1) expression in blood vessels within regions of astrocyte ablation. Mean fluorescence intensity of GLUT1 was measured using ImageJ. As above, Glt1 was used to confirm regions of astrocyte ablation. Images for experimental mice were taken in areas lacking Glt1. 3 confocal images were taken in 3-5 different slices, 3 mice per group. The experimental mice were examined along early (1 dpa), middle (11 dpa) and late (28 dpa) timepoints.

### 2.8 Statistics

Statistics were calculated and graphed using GraphPad Prism 8 (GraphPad Software). Statistics not reported in text can be found in **Table 3**. Data groups were considered significantly different at p≤0.05. Data were tested for Gaussian distribution using the Kolmogorov–Smirnov (KS) normality test. Statistical tests were chosen accordingly and are specified in the results section or figure legends. Data values are reported as mean with standard error of the mean (SEM). Scatter plots reflect individual values and bar graphs reflect the mean with SEM. In graphs with slices plotted, slices from the same animal were plotted as dots of the same color, and different colors represent data points from different mice. Statistical significance is indicated with *p≤0.05, **p≤ 0.01, ***p≤ 0.001, **** p≤ 0.0001.

**Table 3.** Statistics and significance test results

| Figure 1f | Kruskal-Wallis test, significant difference for genotype of animals given tamoxifen and post-administration timepoint (p< 0.001). N is sample number. |  |  |  |  |  |
| --- | --- | --- | --- | --- | --- | --- |
| Dunn's multiple comparisons test | Mean rank diff. | Significant? | Summary | Adjusted P value |  |  |
| Control vs. 2 hpa | -27.54 | No | ns | 0.3851 |  |  |
| Control vs. 6 hpa | -6.536 | No | ns | >0.9999 |  |  |
| Control vs. 1 dpa | -40.98 | Yes | ** | 0.0044 |  |  |
| Control vs. 3 dpa | -47.23 | Yes | *** | 0.0002 |  |  |
| Control vs. 5 dpa | -74.43 | Yes | **** | <0.0001 |  |  |
| Control vs. 11 dpa | -52.05 | Yes | **** | <0.0001 |  |  |
| Control vs. 28 dpa | -49.98 | Yes | *** | 0.0004 |  |  |
| Test details | Mean 1 | SEM 1 | Mean 2 | SEM 2 | N1 | N2 |
| Control vs. 2 hpa | 2.201 | 0.1989 | 4.426 | 0.9110 | 51 | 9 |
| Control vs. 6 hpa | 2.201 | 0.1989 | 2.540 | 0.4053 | 51 | 9 |
| Control vs. 1 dpa | 2.201 | 0.1989 | 6.190 | 1.183 | 51 | 14 |
| Control vs. 3 dpa | 2.201 | 0.1989 | 7.051 | 1.157 | 51 | 16 |
| Control vs. 5 dpa | 2.201 | 0.1989 | 12.620 | 2.026 | 51 | 11 |
| Control vs. 11 dpa | 2.201 | 0.1989 | 8.687 | 1.485 | 51 | 14 |
| Control vs. 28 dpa | 2.201 | 0.1989 | 9.219 | 2.360 | 51 | 13 |
| Figure 1g | Two-way ANOVA, significant difference for genotype of animals given tamoxifen (p<0.0001) and brain region (p=0.0008). N is sample number. |  |  |  |  |  |
| Sidak's multiple comparisons test | Predicted LS Mean Diff. | Significant? | Summary | Adjusted p value |  |  |
| Medial Control vs. Medial Experimental | -7.296 | Yes | **** | <0.0001 |  |  |
| Medial Control vs. Lateral Control | -0.1749 | No | Ns | >0.9999 |  |  |
| Medial Experimental vs. Lateral Experimental | -0.5990 | Yes | **** | <0.0001 |  |  |
| Lateral Control vs. Lateral Experimental | 7.121 | no | ns | 0.9999 |  |  |
| Test Details | Mean 1 | SEM 1 | Mean 2 | SEM 2 | N1 | N2 |
| Medial Control vs. Medial Experimental | 2.436 | 0.246 | 9.732 | 0.863 | 30 | 63 |
| Medial Control vs. Lateral Control | 2.436 | 0.246 | 2.611 | 0.414 | 30 | 22 |
| Medial Experimental vs. Lateral Experimental | 9.732 | 0.863 | 3.035 | 0.579 | 63 | 19 |
| Lateral Control vs. Lateral Experimental | 2.611 | 0.414 | 3.035 | 0.579 | 22 | 19 |
| Figure 2c | One-way ANOVA, significant difference for genotype and post-administration timepoint (p<0.0001). N is sample number. |  |  |  |  |  |
| Tukey's multiple comparisons test | Mean Diff | Significant | Summary | Adjusted p value |  |  |
| Control vs. 2 hpa | -1.188 | No | Ns | 0.9996 |  |  |
| Control vs. 6 hpa | -3.659 | No | Ns | 0.6368 |  |  |
| Control vs. 1 dpa | -7.703 | Yes | ** | 0.0019 |  |  |
| Control vs. 3 dpa | -12.13 | Yes | **** | <0.0001 |  |  |
| Control vs. 5 dpa | -16.84 | Yes | **** | <0.0001 |  |  |
| Control vs. 11 dpa | -16.28 | Yes | **** | <0.0001 |  |  |
| Control vs. 28 dpa | -6.489 | No | Ns | 0.0759 |  |  |
| 11 dpa vs. 28 dpa | 9.791 | Yes | ** | 0.0034 |  |  |
| Test Details | Mean 1 | SEM 1 | Mean 2 | SEM 2 | N1 | N2 |
| Control vs. 2 hpa | 0.1322 | 0.08273 | 1.321 | 0.6469 | 48 | 9 |
| Control vs. 6 hpa | 0.1322 | 0.08273 | 3.792 | 0.9597 | 48 | 12 |
| Control vs. 1 dpa | 0.1322 | 0.08273 | 7.835 | 1.395 | 48 | 15 |
| Control vs. 3 dpa | 0.1322 | 0.08273 | 12.26 | 2.724 | 48 | 13 |
| Control vs. 5 dpa | 0.1322 | 0.08273 | 16.97 | 3.001 | 48 | 12 |
| Control vs. 11 dpa | 0.1322 | 0.08273 | 16.41 | 2.466 | 48 | 19 |
| Control vs. 28 dpa | 0.1322 | 0.08273 | 6.621 | 1.799 | 48 | 10 |
| 11 dpa vs. 28 dpa | 16.41 | 2.466 | 6.621 | 1.799 | 19 | 10 |
| Figure 3b | Kruskal-Wallis test, significant difference for genotype and post-administration timepoint (p<0.0001) |  |  |  |  |  |
| Dunn's multiple comparisons test | Mean rank diff. | Significant? | Summary | Adjusted P value |  |  |
| Control vs. 6 hpa | -2.450 | No | ns | >0.999 |  |  |
| Control vs. 11 dpa | -15.98 | Yes | ** | 0.0011 |  |  |
| Control vs. 28 dpa | -19.39 | Yes | *** | 0.0004 |  |  |
| 6 hpa vs. 11 dpa | -13.53 | No | Ns | 0.0863 |  |  |
| 6 hpa vs. 28 dpa | -16.94 | Yes | * | 0.0285 |  |  |
| 11 dpa vs. 28 dpa | -3.416 | No | Ns | >0.9999 |  |  |
| **Test Details** | **Mean 1** | **SEM 1** | **Mean 2** | **SEM 2** | **N1** | **N2** |
| Control vs. 6 hpa | 2.963 | 0.7270 | 6.731 | 4.021 | 12 | 5 |
| Control vs. 11 dpa | 2.963 | 0.7270 | 28.12 | 3.536 | 12 | 11 |
| Control vs. 28 dpa | 2.963 | 0.7270 | 35.35 | 4.207 | 12 | 7 |
| 6 hpa vs. 11 dpa | 6.731 | 4.021 | 28.12 | 3.536 | 5 | 11 |
| 6 hpa vs. 28 dpa | 6.731 | 4.021 | 35.35 | 4.207 | 5 | 7 |
| 11 dpa vs. 28 dpa | 28.12 | 3.536 | 35.35 | 4.207 | 11 | 7 |
| **Figure 3c** | Kruskal-Wallis test, significant difference for genotype and post-administration timepoint (p=0.0004) |  |  |  |  |  |
| **Dunn's multiple comparisons test** | **Mean rank diff.** | **Significant?** | **Summary** | **Adjusted P value** |  |  |
| Control vs. 6 hpa | 0.6727 | No | Ns | >0.9999 |  |  |
| Control vs. 11 dpa | 12.82 | Yes | * | 0.0152 |  |  |
| Control vs. 28 dpa | 17.13 | Yes | ** | 0.0022 |  |  |
| 6 hpa vs. 11 dpa | 12.15 | No | Ns | 0.1425 |  |  |
| 6 hpa vs. 28 dpa | 16.46 | Yes | * | 0.0286 |  |  |
| 11 dpa vs. 28 dpa | 4.312 | No | Ns | >0.9999 |  |  |
| **Test Details** | **Mean 1** | **SEM 1** | **Mean 2** | **SEM 2** | **N1** | **N2** |
| Control vs. 6 hpa | 1109 | 129 | 1001 | 187 | 11 | 5 |
| Control vs. 11 dpa | 1109 | 129 | 457.4 | 55.81 | 11 | 11 |
| Control vs. 28 dpa | 1109 | 129 | 351.9 | 90.11 | 11 | 7 |
| 6 hpa vs. 11 dpa | 1001 | 187 | 457.4 | 55.81 | 5 | 11 |
| 6 hpa vs. 28 dpa | 1001 | 187 | 351.9 | 90.11 | 5 | 7 |
| 11 dpa vs. 28 dpa | 457.4 | 55.81 | 351.9 | 90.11 | 11 | 7 |
| **Fig 3e** | One-way ANOVA, no significant difference for genotype and post-administration timepoint (p=0.9960) |  |  |  |  |  |
| **Dunnett's multiple comparisons test** | **Mean Diff** | **Significant?** | **Summary** | **Adjusted P value** |  |  |
| Control vs. 6 hpa | -11.85 | No | Ns | 0.9996 |  |  |
| Control vs. 11 dpa | -31.08 | No | Ns | 0.9946 |  |  |
| Control vs. 28 dpa | -33.26 | No | Ns | 0.9934 |  |  |
| **Test Details** | **Mean 1** | **SEM 1** | **Mean 2** | **SEM 2** | **N1** | **N2** |
| Control vs. 6 hpa | 423.4 | 171.7 | 435.2 | 106.5 | 3 | 3 |
| Control vs. 11 dpa | 423.4 | 171.7 | 454.5 | 29.52 | 3 | 3 |
| Control vs. 28 dpa | 423.4 | 171.7 | 456.7 | 101.3 | 3 | 3 |
| **Figure 5b** | One-Way ANOVA, significant difference for post-administration timepoint (p= 0.0273), N is sample number. |  |  |  |  |  |
| **Tukey's multiple comparisons test** | **Significant?** | **Summary** | **Adjusted P value** |  |  |  |
| 1 dpa vs. 11 dpa | No | ns | 0.6083 |  |  |  |
| 1 dpa vs. 28 dpa | Yes | * | 0.0228 |  |  |  |
| 11 dpa vs. 28 dpa | No | ns | 0.1810 |  |  |  |
| **Test details** | **Mean 1** | **Mean 2** | **Mean Diff.** | **SE of diff.** | **N1** | **N2** |
| 1 dpa vs. 11 dpa | 815.9 | 938.8 | -122.9 | 128.5 | 15 | 15 |
| 1 dpa vs. 28 dpa | 815.9 | 1171 | -354.6 | 128.5 | 15 | 15 |
| 11 dpa vs. 28 dpa | 938.8 | 1171 | -231.7 | 128.5 | 15 | 15 |
| **Figure 5c** | One-Way ANOVA, significant difference for post-administration timepoint ($p = 0.0013$), N is sample number. | | | | | |
| **Tukey's multiple comparisons test** | **Significant?** | **Summary** | **Adjusted P value** |  |  |  |
| 1 dpa vs. 11 dpa | No | ns | 0.6446 |  |  |  |
| 1 dpa vs. 28 dpa | Yes | * | 0.0204 |  |  |  |
| 11 dpa vs. 28 dpa | No | ns | 0.1499 |  |  |  |
| **Test details** | **Mean 1** | **Mean 2** | **Mean Diff.** | **SE of diff.** | **N1** | **N2** |
| 1 dpa vs. 11 dpa | 1086 | 1402 | -316.2 | 352.2 | 15 | 15 |
| 1 dpa vs. 28 dpa | 1086 | 2073 | -987.0 | 352.2 | 15 | 15 |
| 11 dpa vs. 28 dpa | 1402 | 2073 | -670.8 | 352.2 | 15 | 15 |
| **Figure 5d** | Two-way ANOVA significance, significant difference for post-administration timepoint ($p < 0.0001$) and intersection distance ($p < 0.0001$). N is sample number. | | | | | |
| **Uncorrected Fisher's LSD test** | **Predicted Mean Diff.** | **Significant?** | **Summary** | **Adjusted P value** |  |  |
| **Intersection distance 1** |  |  |  |  |  |  |
| 1 dpa vs. 11 dpa | -0.2 | No | ns | 0.9341 |  |  |
| 1 dpa vs. 28 dpa | -0.5333 | No | ns | 0.8256 |  |  |
| 11 dpa vs. 28 dpa | -0.3333 | No | ns | 0.8904 |  |  |
| **Intersection distance 2** | **Predicted Mean Diff.** | **Significant?** | **Summary** | **Adjusted P value** |  |  |
| 1 dpa vs. 11 dpa | -2.733 | No | ns | 0.2588 |  |  |
| 1 dpa vs. 28 dpa | -0.4667 | No | ns | 0.8471 |  |  |
| 11 dpa vs. 28 dpa | 2.267 | No | ns | 0.349 |  |  |
| **Intersection distance 3** | **Predicted Mean Diff.** | **Significant?** | **Summary** | **Adjusted P value** |  |  |
| 1 dpa vs. 11 dpa | 0.6667 | No | ns | 0.7829 |  |  |
| 1 dpa vs. 28 dpa | 0.8667 | No | ns | 0.7202 |  |  |
| 11 dpa vs. 28 dpa | 0.2 | No | ns | 0.9341 |  |  |
| **Intersection distance 4** | **Predicted Mean Diff.** | **Significant?** | **Summary** | **Adjusted P value** |  |  |
| 1 dpa vs. 11 dpa | -2.533 | No | ns | 0.2953 |  |  |
| 1 dpa vs. 28 dpa | 0.2667 | No | ns | 0.9122 |  |  |
| 11 dpa vs. 28 dpa | 2.8 | No | ns | 0.2474 |  |  |
| **Intersection distance 5** | **Predicted Mean Diff.** | **Significant?** | **Summary** | **Adjusted P value** |  |  |
| 1 dpa vs. 11 dpa | -2.667 | No | ns | 0.2706 |  |  |
| 1 dpa vs. 28 dpa | 0.6667 | No | ns | 0.7829 |  |  |
| 11 dpa vs. 28 dpa | 3.333 | No | ns | 0.1686 |  |  |
| **Intersection distance 6** | **Predicted Mean Diff.** | **Significant?** | **Summary** | **Adjusted P value** |  |  |
| 1 dpa vs. 11 dpa | -0.5333 | No | ns | 0.8256 |  |  |
| 1 dpa vs. 28 dpa | -0.8 | No | ns | 0.7409 |  |  |
| 11 dpa vs. 28 dpa | -0.2667 | No | ns | 0.9122 |  |  |
| **Intersection distance 7** | **Predicted Mean Diff.** | **Significant?** | **Summary** | **Adjusted P value** |  |  |
| 1 dpa vs. 11 dpa | -2.933 | No | ns | 0.2256 |  |  |
| 1 dpa vs. 28 dpa | -5.467 | Yes | * | 0.0241 |  |  |
| 11 dpa vs. 28 dpa | -2.533 | No | ns | 0.2953 |  |  |
| **Intersection distance 8** | **Predicted Mean Diff.** | **Significant?** | **Summary** | **Adjusted P value** |  |  |
| 1 dpa vs. 11 dpa | 0 | No | ns | >0.9999 |  |  |
| 1 dpa vs. 28 dpa | -5.667 | Yes | * | 0.0194 |  |  |
| 11 dpa vs. 28 dpa | -5.667 | Yes | * | 0.0194 |  |  |
| **Intersection distance 9** | **Predicted Mean Diff.** | **Significant?** | **Summary** | **Adjusted P value** |  |  |
| 1 dpa vs. 11 dpa | -2.067 | No | ns | 0.3932 |  |  |
| 1 dpa vs. 28 dpa | -7.2 | Yes | ** | 0.003 |  |  |
| 11 dpa vs. 28 dpa | -5.133 | Yes | * | 0.0341 |  |  |
| **Intersection distance 10** | **Predicted Mean Diff.** | **Significant?** | **Summary** | **Adjusted P value** |  |  |
| 1 dpa vs. 11 dpa | -1.6 | No | ns | 0.5085 |  |  |
| 1 dpa vs. 28 dpa | -6.2 | Yes | * | 0.0106 |  |  |
| 11 dpa vs. 28 dpa | -4.6 | No | ns | 0.0576 |  |  |
| **Intersection distance 11** | **Predicted Mean Diff.** | **Significant?** | **Summary** | **Adjusted P value** |  |  |
| 1 dpa vs. 11 dpa | -0.8 | No | ns | 0.7409 |  |  |
| 1 dpa vs. 28 dpa | -6.267 | Yes | ** | 0.0098 |  |  |
| 11 dpa vs. 28 dpa | -5.467 | Yes | * | 0.0241 |  |  |
| **Intersection distance 12** | **Predicted Mean Diff.** | **Significant?** | **Summary** | **Adjusted P value** |  |  |
| 1 dpa vs. 11 dpa | -0.06667 | No | ns | 0.978 |  |  |
| 1 dpa vs. 28 dpa | -5.533 | Yes | * | 0.0224 |  |  |
| 11 dpa vs. 28 dpa | -5.467 | Yes | * | 0.0241 |  |  |
| **Intersection distance 13** | **Predicted Mean Diff.** | **Significant?** | **Summary** | **Adjusted P value** |  |  |
| 1 dpa vs. 11 dpa | -0.3857 | No | ns | 0.8755 |  |  |
| 1 dpa vs. 28 dpa | -5.452 | Yes | * | 0.0271 |  |  |
| 11 dpa vs. 28 dpa | -5.067 | Yes | * | 0.0365 |  |  |
| **Intersection distance 14** | **Predicted Mean Diff.** | **Significant?** | **Summary** | **Adjusted P value** |  |  |
| 1 dpa vs. 11 dpa | -0.3333 | No | ns | 0.8904 |  |  |
| 1 dpa vs. 28 dpa | -3.8 | No | ns | 0.1166 |  |  |
| 11 dpa vs. 28 dpa | -3.467 | No | ns | 0.1522 |  |  |
| **Intersection distance 15** | **Predicted Mean Diff.** | **Significant?** | **Summary** | **Adjusted P value** |  |  |
| 1 dpa vs. 11 dpa | -0.06667 | No | ns | 0.978 |  |  |
| 1 dpa vs. 28 dpa | -1.6 | No | ns | 0.5085 |  |  |
| 11 dpa vs. 28 dpa | -1.533 | No | ns | 0.5263 |  |  |
| **Intersection distance 16** | **Predicted Mean Diff.** | **Significant?** | **Summary** | **Adjusted P value** |  |  |
| 1 dpa vs. 11 dpa | 0 | No | ns | >0.9999 |  |  |
| 1 dpa vs. 28 dpa | -0.9333 | No | ns | 0.6997 |  |  |
| 11 dpa vs. 28 dpa | -0.9333 | No | ns | 0.6997 |  |  |
| **Intersection distance 17** | **Predicted Mean Diff.** | **Significant?** | **Summary** | **Adjusted P value** |  |  |
| 1 dpa vs. 11 dpa | 0 | No | ns | >0.9999 |  |  |
| 1 dpa vs. 28 dpa | -0.3333 | No | ns | 0.8904 |  |  |
| 11 dpa vs. 28 dpa | -0.3333 | No | ns | 0.8904 |  |  |
| **Intersection distance 18** | **Predicted Mean Diff.** | **Significant?** | **Summary** | **Adjusted P value** |  |  |
| 1 dpa vs. 11 dpa | 0 | No | ns | >0.9999 |  |  |
| 1 dpa vs. 28 dpa | -0.2 | No | ns | 0.9341 |  |  |
| 11 dpa vs. 28 dpa | -0.2 | No | ns | 0.9341 |  |  |
| **Test Details** | **Mean 1** | **SEM 1** | **Mean 2** | **SEM 2** | **N1** | **N2** |
| **Intersection distance 1** |  |  |  |  |  |  |
| 1 dpa vs. 11 dpa | 2.2000 | 0.1745 | 2.4000 | 0.2894 | 15 | 15 |
| 1 dpa vs. 28 dpa | 2.2000 | 0.1745 | 2.7333 | 0.4414 | 15 | 15 |
| 11 dpa vs. 28 dpa | 2.4000 | 0.2894 | 2.7333 | 0.4414 | 15 | 15 |
| **Intersection distance 2** | **Mean 1** | **SEM 1** | **Mean 2** | **SEM 2** | **N1** | **N2** |
| 1 dpa vs. 11 dpa | 6.3333 | 0.7411 | 9.0667 | 1.0257 | 15 | 15 |
| 1 dpa vs. 28 dpa | 6.3333 | 0.7411 | 6.800 | 0.8519 | 15 | 15 |
| 11 dpa vs. 28 dpa | 9.0667 | 1.0257 | 6.800 | 0.8519 | 15 | 15 |
| **Intersection distance 3** | **Mean 1** | **SEM 1** | **Mean 2** | **SEM 2** | **N1** | **N2** |
| 1 dpa vs. 11 dpa | 12.8000 | 1.8752 | 12.1333 | 0.9354 | 15 | 15 |
| 1 dpa vs. 28 dpa | 12.8000 | 1.8752 | 11.9333 | 1.5568 | 15 | 15 |
| 11 dpa vs. 28 dpa | 12.1333 | 0.9354 | 11.9333 | 1.5568 | 15 | 15 |
| **Intersection distance 4** | **Mean 1** | **SEM 1** | **Mean 2** | **SEM 2** | **N1** | **N2** |
| 1 dpa vs. 11 dpa | 16.7333 | 1.8605 | 19.2667 | 1.4879 | 15 | 15 |
| 1 dpa vs. 28 dpa | 16.7333 | 1.8605 | 16.4667 | 2.2988 | 15 | 15 |
| 11 dpa vs. 28 dpa | 19.2667 | 1.4879 | 16.4667 | 2.2988 | 15 | 15 |
| **Intersection distance 5** | **Mean 1** | **SEM 1** | **Mean 2** | **SEM 2** | **N1** | **N2** |
| 1 dpa vs. 11 dpa | 21.1333 | 1.9417 | 23.8000 | 2.0775 | 15 | 15 |
| 1 dpa vs. 28 dpa | 21.1333 | 1.9417 | 20.4667 | 2.6147 | 15 | 15 |
| 11 dpa vs. 28 dpa | 23.8000 | 2.0775 | 20.4667 | 2.6147 | 15 | 15 |
| **Intersection distance 6** | **Mean 1** | **SEM 1** | **Mean 2** | **SEM 2** | **N1** | **N2** |
| 1 dpa vs. 11 dpa | 22.0667 | 2.2980 | 22.6000 | 2.5219 | 15 | 15 |
| 1 dpa vs. 28 dpa | 22.0667 | 2.2980 | 22.8667 | 3.5144 | 15 | 15 |
| 11 dpa vs. 28 dpa | 22.6000 | 2.5219 | 22.8667 | 3.5144 | 15 | 15 |
| **Intersection distance 7** | **Mean 1** | **SEM 1** | **Mean 2** | **SEM 2** | **N1** | **N2** |
| 1 dpa vs. 11 dpa | 18.9333 | 2.5699 | 21.8667 | 2.5799 | 15 | 15 |
| 1 dpa vs. 28 dpa | 18.9333 | 2.5699 | 24.4000 | 3.7299 | 15 | 15 |
| 11 dpa vs. 28 dpa | 21.8667 | 2.5799 | 24.4000 | 3.7299 | 15 | 15 |
| **Intersection distance 8** | **Mean 1** | **SEM 1** | **Mean 2** | **SEM 2** | **N1** | **N2** |
| 1 dpa vs. 11 dpa | 14.4000 | 2.3559 | 14.4000 | 1.9019 | 15 | 15 |
| 1 dpa vs. 28 dpa | 14.4000 | 2.3559 | 20.0667 | 3.2870 | 15 | 15 |
| 11 dpa vs. 28 dpa | 14.4000 | 1.9019 | 20.0667 | 3.2870 | 15 | 15 |
| **Intersection distance 9** | **Mean 1** | **SEM 1** | **Mean 2** | **SEM 2** | **N1** | **N2** |
| 1 dpa vs. 11 dpa | 7.8000 | 1.5681 | 9.8667 | 1.7206 | 15 | 15 |
| 1 dpa vs. 28 dpa | 7.8000 | 1.5681 | 15.0000 | 2.4708 | 15 | 15 |
| 11 dpa vs. 28 dpa | 9.8667 | 1.7206 | 15.0000 | 2.4708 | 15 | 15 |
| **Intersection distance 10** | **Mean 1** | **SEM 1** | **Mean 2** | **SEM 2** | **N1** | **N2** |
| 1 dpa vs. 11 dpa | 4.4667 | 1.4305 | 6.0667 | 1.3504 | 15 | 15 |
| 1 dpa vs. 28 dpa | 4.4667 | 1.4305 | 10.6667 | 2.6034 | 15 | 15 |
| 11 dpa vs. 28 dpa | 6.0667 | 1.3504 | 10.6667 | 2.6034 | 15 | 15 |
| **Intersection distance 11** | **Mean 1** | **SEM 1** | **Mean 2** | **SEM 2** | **N1** | **N2** |
| 1 dpa vs. 11 dpa | 2.3333 | 1.1819 | 3.1333 | 1.2223 | 15 | 15 |
| 1 dpa vs. 28 dpa | 2.3333 | 1.1819 | 8.6000 | 2.3620 | 15 | 15 |
| 11 dpa vs. 28 dpa | 3.1333 | 1.2223 | 8.6000 | 2.3620 | 15 | 15 |
| **Intersection distance 12** | **Mean 1** | **SEM 1** | **Mean 2** | **SEM 2** | **N1** | **N2** |
| 1 dpa vs. 11 dpa | 1.0000 | 0.4781 | 1.0667 | 0.5297 | 15 | 15 |
| 1 dpa vs. 28 dpa | 1.0000 | 0.4781 | 6.5333 | 2.3822 | 15 | 15 |
| 11 dpa vs. 28 dpa | 1.0667 | 0.5297 | 6.5333 | 2.3822 | 15 | 15 |
| **Intersection distance 13** | **Mean 1** | **SEM 1** | **Mean 2** | **SEM 2** | **N1** | **N2** |
| 1 dpa vs. 11 dpa | 0.2143 | 0.1547 | 0.6000 | 0.3754 | 15 | 15 |
| 1 dpa vs. 28 dpa | 0.2143 | 0.1547 | 5.6667 | 2.7372 | 15 | 15 |
| 11 dpa vs. 28 dpa | 0.6000 | 0.3754 | 5.6667 | 2.7372 | 15 | 15 |
| **Intersection distance 14** | **Mean 1** | **SEM 1** | **Mean 2** | **SEM 2** | **N1** | **N2** |
| 1 dpa vs. 11 dpa | 0.0000 | 0.0000 | 0.3333 | 0.3333 | 15 | 15 |
| 1 dpa vs. 28 dpa | 0.0000 | 0.0000 | 3.8000 | 2.2215 | 15 | 15 |
| 11 dpa vs. 28 dpa | 0.3333 | 0.3333 | 3.8000 | 2.2215 | 15 | 15 |
| **Intersection distance 15** | **Mean 1** | **SEM 1** | **Mean 2** | **SEM 2** | **N1** | **N2** |
| 1 dpa vs. 11 dpa | 0.0000 | 0.0000 | 0.0667 | 0.0667 | 15 | 15 |
| 1 dpa vs. 28 dpa | 0.0000 | 0.0000 | 1.6000 | 1.2024 | 15 | 15 |
| 11 dpa vs. 28 dpa | 0.0667 | 0.0667 | 1.6000 | 1.2024 | 15 | 15 |
| **Intersection distance 16** | **Mean 1** | **SEM 1** | **Mean 2** | **SEM 2** | **N1** | **N2** |
| 1 dpa vs. 11 dpa | 0.0000 | 0.0000 | 0.0000 | 0.0000 | 15 | 15 |
| 1 dpa vs. 28 dpa | 0.0000 | 0.0000 | 0.9333 | 0.5206 | 15 | 15 |
| 11 dpa vs. 28 dpa | 0.0000 | 0.0000 | 0.9333 | 0.5206 | 15 | 15 |
| **Intersection distance 17** | **Mean 1** | **SEM 1** | **Mean 2** | **SEM 2** | **N1** | **N2** |
| 1 dpa vs. 11 dpa | 0.0000 | 0.0000 | 0.0000 | 0.0000 | 15 | 15 |
| 1 dpa vs. 28 dpa | 0.0000 | 0.0000 | 0.3330 | 0.1869 | 15 | 15 |
| 11 dpa vs. 28 dpa | 0.0000 | 0.0000 | 0.3330 | 0.1869 | 15 | 15 |
| **Intersection distance 18** | **Mean 1** | **SEM 1** | **Mean 2** | **SEM 2** | **N1** | **N2** |
| 1 dpa vs. 11 dpa | 0.0000 | 0.0000 | 0.0000 | 0.0000 | 15 | 15 |
| 1 dpa vs. 28 dpa | 0.0000 | 0.0000 | 0.2000 | 0.1447 | 15 | 15 |
| 11 dpa vs. 28 dpa | 0.0000 | 0.0000 | 0.2000 | 0.1447 | 15 | 15 |
| **Figure 5f** | One-way ANOVA, no significant difference for genotype and post-administration timepoint (p=0.1382) |  |  |  |  |  |
| **Tukey's Multiple Comparisons test** | **Mean Diff** | **Significant?** | **Summary** | **Adjusted P value** |  |  |
| Control vs. 1 dpa | 0.1601 | No | ns | 0.2119 |  |  |
| Control vs. 11 dpa | 0.1576 | No | ns | 0.3058 |  |  |
| Control vs. 28 dpa | 0.1199 | No | ns | 0.4944 |  |  |
| 1 dpa vs. 11 dpa | -0.002496 | No | ns | >0.9999 |  |  |
| 1 dpa vs. 28 dpa | -0.04018 | No | ns | 0.9758 |  |  |
| 11 dpa vs. 28 dpa | -0.03768 | No | ns | 0.9836 |  |  |

| Test Details | Mean 1 | SEM 1 | Mean 2 | SEM 2 | N1 | N2 |
| --- | --- | --- | --- | --- | --- | --- |
| Control vs. 1 dpa | 1.005 | 0.0429 | 0.8447 | 0.06033 | 17 | 8 |
| Control vs. 11 dpa | 1.005 | 0.0429 | 0.8472 | 0.0732 | 17 | 6 |
| Control vs. 28 dpa | 1.005 | 0.0429 | 0.8849 | 0.08895 | 17 | 7 |
| 1 dpa vs. 11 dpa | 0.8447 | 0.06033 | 0.8472 | 0.0732 | 8 | 6 |
| 1 dpa vs. 28 dpa | 0.8447 | 0.06033 | 0.8849 | 0.08895 | 8 | 7 |
| 11 dpa vs. 28 dpa | 0.8472 | 0.0732 | 0.8849 | 0.08895 | 6 | 7 |
| **Figure 5g** | One-way ANOVA, no significant difference for genotype and post-administration timepoint (p=0.1896) |  |  |  |  |  |
| Tukey's Multiple Comparisons test | Mean Diff | Significant? | Summary | Adjusted P value |  |  |
| Control vs. 1 dpa | 0.2294 | No | ns | 0.1712 |  |  |
| Control vs. 11 dpa | -0.007537 | No | ns | >0.9999 |  |  |
| Control vs. 28 dpa | 0.09268 | No | ns | 0.8479 |  |  |
| 1 dpa vs. 11 dpa | -0.2369 | No | ns | 0.3253 |  |  |
| 1 dpa vs. 28 dpa | -0.1367 | No | ns | 0.7272 |  |  |
| 11 dpa vs. 28 dpa | 0.1002 | No | ns | 0.8926 |  |  |
| Test Details | Mean 1 | SEM 1 | Mean 2 | SEM 2 | N1 | N2 |
| Control vs. 1 dpa | 1.002 | 0.06148 | 0.7728 | 0.09323 | 17 | 8 |
| Control vs. 11 dpa | 1.002 | 0.06148 | 1.01 | 0.1398 | 17 | 6 |
| Control vs. 28 dpa | 1.002 | 0.06148 | 0.9095 | 0.04635 | 17 | 7 |
| 1 dpa vs. 11 dpa | 0.7728 | 0.09323 | 1.01 | 0.1398 | 8 | 6 |
| 1 dpa vs. 28 dpa | 0.7728 | 0.09323 | 0.9095 | 0.04635 | 8 | 7 |
| 11 dpa vs. 28 dpa | 1.01 | 0.1398 | 0.9095 | 0.04635 | 6 | 7 |

## 3. Results

### 3.1 Astrocyte ablation occurs within hours after Tamoxifen administration

To determine if astrocytes are necessary for blood-brain barrier (BBB) maintenance, we genetically ablated a small number of astrocytes in adult mice using a conditional and inducible approach. Mice expressing the diphtheria toxin subunit A (DTA) behind a stop cassette flanked by loxP sites were bred with mice expressing CreERT behind the promoter of the astrocytic glutamate transporter Glast. This restricted expression of CreERT to astrocytes and enabled induction of astrocyte ablation in adult mice upon administration of Tamoxifen, which causes translocation of the Cre recombinase to the nucleus. There, Cre recombinase excised the stop cassette enabling expression of DTA (**Fig. 1a**). We chose Glast-CreERT specifically because of its limited recombination efficiency in the adult forebrain (Mori et al., 2006), which allowed for sparse astrocyte ablation and confirmed that no astrocyte ablation took place in experimental DTA^fl/wt^//Glast-CreERT^tg/wt^ mice in the absence of Tamoxifen administration (**Suppl. Fig. 2a**).

We first determined the time course of astrocyte ablation by harvesting brain tissue 2h, 6h, 24h, 3d, 5d, 11d and 28d after a single dose of Tamoxifen (Tx) or carrier solution (10% ethanol in corn oil) (**Fig. 1b**). At 6 hpa and 1 dpa, we observed overlap between the astrocyte marker S100β and the apoptosis marker cleaved caspase 3, suggesting that astrocytes underwent apoptosis at these timepoints **(Fig. 1c)**. The nuclei in these astrocytes were pyknotic and highly condensed, an additional indication that these cells were undergoing apoptosis. We did not find any cleaved caspase 3-positive cells at later timepoints (3 dpa, 5 dpa, 11 dpa, 28 dpa), suggesting that by those timepoints, the recombined astrocytes had died.

We confirmed astrocyte loss across all timepoints using the astrocyte membrane-associated protein glutamate transporter 1 (Glt1). Tissues from control mice were characterized by a mostly even and continuous Glt1 staining pattern (**Fig. 1d**), though astrocytes close to the midline sometimes had reduced Glt1 levels in both control and experimental groups (**Fig. 1e**). After tamoxifen administration DTA^fl/wt^//Glast-CreERT^tg/wt^ mice had disrupted Glt1 staining patterns in distinct regions resembling single astrocyte domains or groups of astrocytes throughout the cortical gray matter (**Fig. 1d-e**) and other brain regions, including the hippocampus, striatum and cerebellum (**Suppl. Fig. 2b-d**).

Areas lacking Glt1 were also characterized by lack of expression of other astrocyte proteins including S100β, and Aquaporin4 (AQ4) (**Suppl. Fig. 3a-b**). This occurred, in single domains, as early as 2 hours post ablation (hpa, refers to administration of Tx) (**Fig. 1d-e**). At this timepoint we did not detect cleaved caspase-3 possibly due to the small number of apoptotic astrocytes at this early timepoint.

Quantification of Glt1-negative area size was not sensitive enough to detect significant differences between control and experimental mice at 2 hpa and 6 hpa (**Fig. 1f**), even though small areas with missing astrocytes were clearly present, even at these early timepoints. Cortical area size lacking Glt1 increased at 1 dpa and was maximal at approximately 12% at 5 dpa. At 11 and 28 dpa, areas without Glt1 appeared slightly reduced in size when compared to 5 dpa, but this difference was not statistically significant. For each experimental animal, the largest cortical Glt1 loss areas were consistently located closer to the midline, while smaller loss areas were found primarily in lateral sections. When these regions were plotted separately, medial slices (Allen Brain Atlas slices 8-21) showed significantly larger Glt1 loss areas compared to lateral slices (Allen Brain Atlas slices 1-7) in experimental but not in control mice **(Fig. 1g).**

In conclusion, astrocyte ablation was initiated in DTA^fl/wt^//Glast-CreERT^tg/wt^ mice within hours after Tx administration and peaked at 5 dpa. Approximately 12% of the cortical surface area lacked Glt1 coverage.

### 3.2 Astrocyte ablation triggers early and sustained blood-brain barrier dysfunction

It is unclear if pericytes are sufficient to maintain the adult BBB or if astrocytes are indeed necessary, possibly upstream of pericytes, to maintain BBB function *in vivo*. To determine whether genetic ablation of astrocytes in the adult mouse brain interferes with BBB integrity, we used the small fluorescently labeled molecule Cadaverine (<1kDa). We chose Cadaverine as opposed to larger molecular size tracers as a starting point to detect even small disturbances in BBB function. A recent study reported Cadaverine leakage into the brain parenchyma to be indicative of impaired tight junctions even in the absence of leakage of larger plasma proteins (∼70kDa) or Dextran-coupled fluorescent tracers (10kDa) (Yanagida et al., 2017).

Cadaverine leakage was observed as early as 2 hpa and was typically associated with areas of astrocyte ablation **(Fig. 2a)** suggesting that BBB integrity was affected very quickly once astrocytes become dysfunctional as a result of DTA expression, which inhibits protein synthesis ultimately leading to cell death. Cadaverine was injected 30 minutes before mice were euthanized, thus leakage of the tracer represents the dysfunction of the barrier at this particular time point.

**Figure 2.**
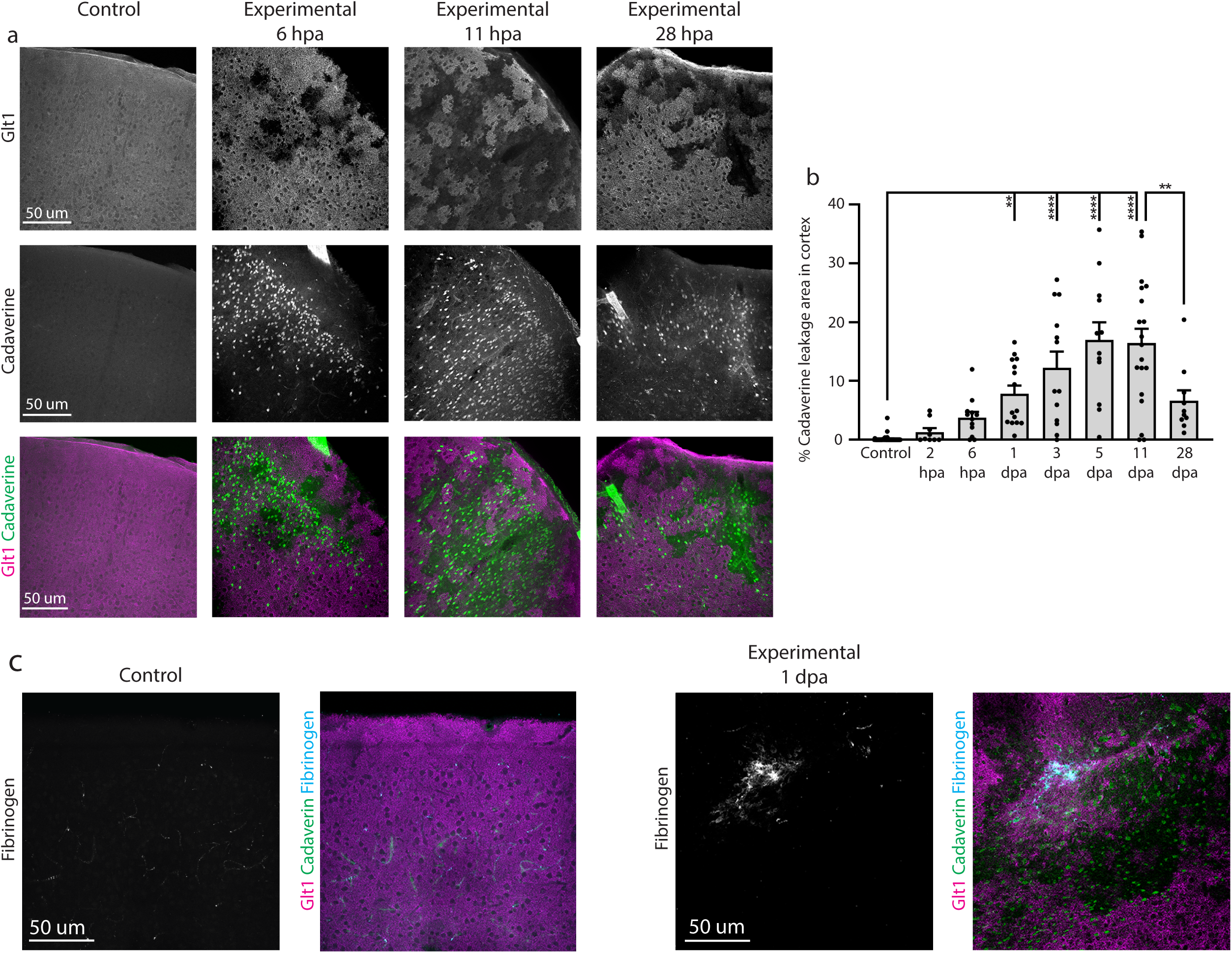
Astrocyte ablation induced BBB dysfunction. **a.** Leakage of the BBB tracer dye Cadaverine in cortex was observed around regions of astrocyte cell death indicated by lack of Glt1 expression. Cadaverine leakage occurred as early as 2 hpa and was still present at 28 dpa. Control mice showed minimal leakage of dye. **b.** Cadaverine leakage occurred across cortex in experimental mice and was quantified as percent leakage area out of entire cortex area and plotted by slice. **c.** Some vessels labeled positive for fibrinogen deposition and were surrounded by Glt1 loss and Cadaverine leakage.

As an approximation of the extent of BBB dysfunction after astrocyte ablation, we quantified the percentage of cortical area covered by cells that took up Cadaverine in large image scans (**Fig. 2b, Suppl. Fig. 4a)**. At 6 hpa about 5% of the cortical gray matter was covered by Cadaverine^+^ cells. This area fraction tripled by 5-11dpa. While Cadaverine spread throughout the tissue appeared diffuse at early timepoints after ablation (6 hpa, 1 dpa) it was found to be more restricted to areas of ablation one month after ablation (28 dpa) signified by a significantly reduced area covered by Cadaverine^+^ cells. We also occasionally found some Cadaverine leakage, albeit at much smaller extent, in control mice. Interestingly, leakages colocalized with areas along the midline where we found astrocytes with reduced Glt1 levels even in controls. Sometimes leakage of Cadaverine occurred in controls surrounding large penetrating arteries with adjacent astrocytes lacking Glt1.

We next sought to determine if greater Cadaverine leakage correlated with greater Glt1 loss by plotting these values from all experimental and control mice. The r value of 0.71 (two-tailed Pearson’s correlation test) reflects a moderate correlation but suggests that Cadaverine leakage is not necessarily larger in mice with larger Glt1 loss regions. This might be due to the small size of the dye, which might easily diffuse within the tissue.

To further determine the extent and severity of BBB dysfunction after astrocyte ablation, we stained for the large molecular size blood plasma protein fibrinogen (∼340 kDa). Fibrinogen accumulates outside of blood vessels and acts as part of the coagulation cascade that stops bleeding, and is indicative of damage to blood vessels resulting in entrance of larger blood-borne molecules and erythrocytes. While not as widespread as Cadaverine leakage, we observed several regions with fibrinogen deposition in the cortex as early as 1 dpa (**Fig. 2c**). In the 14 experimental mice examined for fibrinogen, 7 of the 80 slices examined showed fibrinogen deposition. These regions were consistently surrounded by large zones depleted of Glt1^+^ astrocytes and Cadaverine leakage. However, most areas with Glt1 loss and Cadaverine leakage lacked fibrinogen deposits.

Taken together, these findings suggest that astrocyte ablation interferes with BBB integrity, that larger areas of loss result in more extensive damage to the BBB, allowing entrance of molecules as large as fibrinogen into the brain, and that BBB integrity is disturbed for at least 4 weeks post ablation.

### 3.3 Astrocyte ablation interferes with tight junction function but does not affect expression of endothelial glucose transporter Glut1

We next asked whether Cadaverine and fibrinogen leakage occurred due to dysfunctional tight junctions in astrocyte ablated areas. We examined the tight junction protein zonula occludens-1 (ZO-1) using immunohistochemistry at 6 hpa, 11 dpa and 28 dpa. In control mice, we observed continuous labeling of ZO-1 that overlapped with the endothelial cell marker CD31 (**Fig. 3a**). We observed mostly intact ZO-1 labeling in ablation regions at 6 hpa. These areas did not show Cadaverine leakage. In contrast, the few areas that had disrupted ZO-1 labeling also presented with Cadaverine leakage (**Fig. 3a**). We observed frequent ZO-1 disruption and reduced expression at 11 dpa and 28 dpa, and some vessels lacked ZO-1 almost completely (**Fig. 3a,** yellow arrowhead). Quantifications for ZO-1 intensity and continuity along CD31 vessels at 6 hpa, when we observed only few missing astrocytes domains, showed no changes from controls (**Fig. 3b-c**). At 11 dpa and 28 dpa, the percentage of CD31 vessels lacking ZO-1 labeling was increased and ZO-1 signal intensity was decreased.

**Figure 3.**
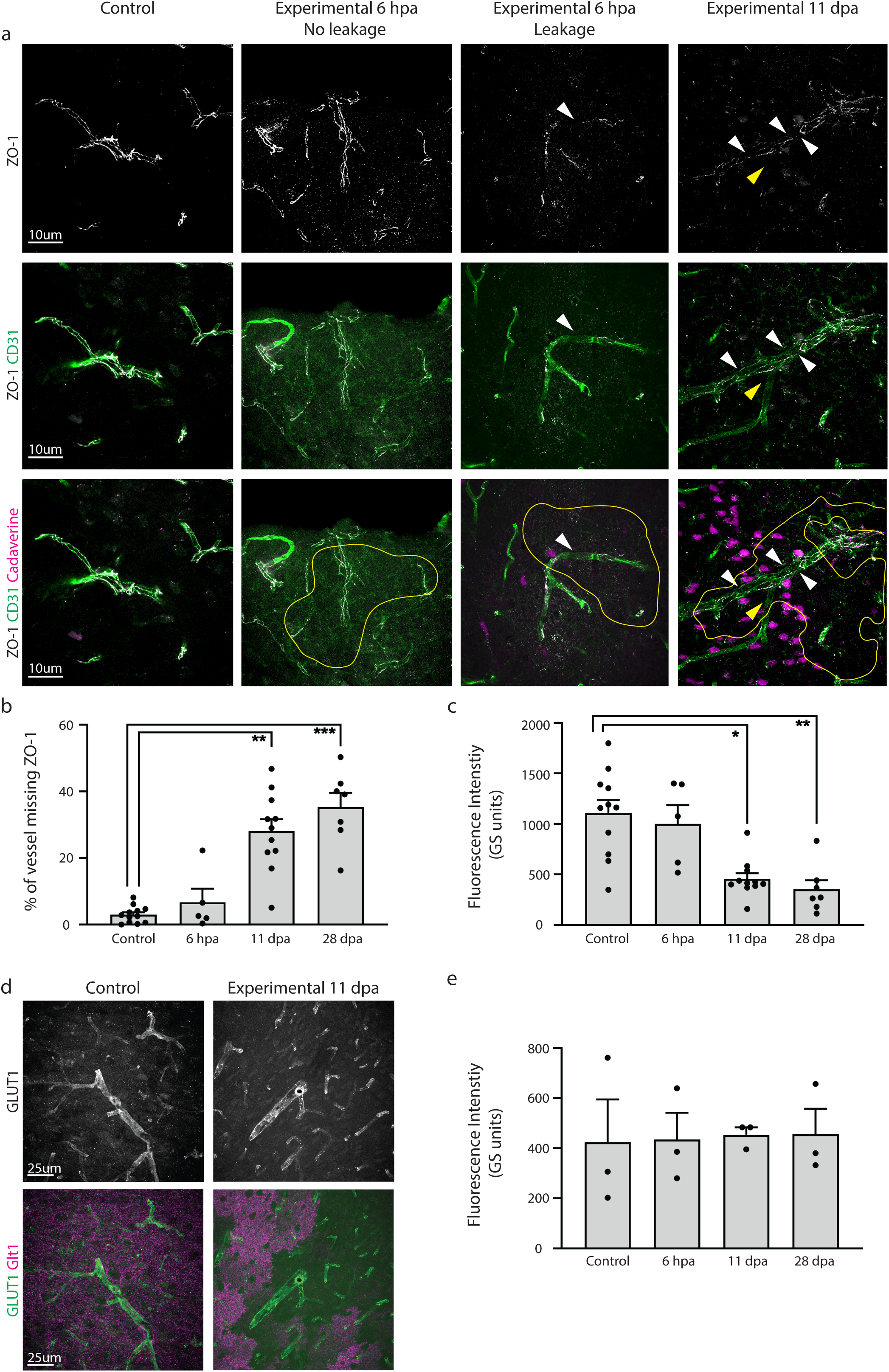
Astrocyte ablation of cells adjacent to blood vessels reduced expression of proteins in endothelial cells responsible for maintenance of the BBB. **a.** Astrocyte ablation of cells next to blood vessels resulted in reduced and discontinuous vessel labeling for the tight junction protein ZO-1 (white arrowheads). ZO-1 expression was examined within Glt-1-lacking areas (circled in yellow). At 6 hpa, few vessels within Glt1-lacking areas showed disrupted ZO-1 labeling, and those that did also presented with Cadaverine leakage. At 11 dpa, some vessels in regions of ablation completely lacked ZO-1 (yellow arrowhead) and showed Cadaverine leakage. **b.** Continuity of ZO-1 labeling in vessels was quantified by binarizing ZO-1 signal and drawing a line along ZO-1 signal, using CD31 as a guide. **c.** Average fluorescence intensity was quantified for ZO-1 intensity profile measurements of lines drawn along ZO-1 labeling. **d.** Endothelial glucose transporter GLUT1 expression was unchanged between groups. **e.** Quantification of average fluorescence intensity of GLUT1 showed no changes in vessels adjacent to Glt1-lacking areas.

Increased permeability due to BBB dysfunction can occur through several mechanisms beyond loosening of physical tightness of the junctions, including dysregulation of proteins that govern its metabolic barrier properties. The endothelial glucose transporter Glut1 is responsible for over 90% of all glucose transport into the brain, and is thus often used as a readout for metabolic barrier function (Boado & Pardridge, 1993; Pardridge, Boado, & Farrell, 1990). We labeled blood vessels with the glucose transporter Glut1 to determine if glucose transporter levels were affected by astrocyte ablation. We found no changes in Glut1 expression pattern at 6 hpa, 11 dpa or 28 dpa **(Fig. 3d)**. Quantification of average fluorescence intensity of Glut1 in vessels adjacent to astrocyte ablation regions revealed no differences among groups (**Fig. 3e).**

Together, this suggests that astrocyte ablation interferes with specific aspects of barrier function such as expression and/or localization of tight junction proteins.

### 3.4 Astrocytes adjacent to ablated areas respond with molecular changes characteristic for glial scar formation

In response to injury and disease, astrocytes undergo context-dependent molecular and morphological changes in a process called astrogliosis. In the event of a focal traumatic brain or spinal cord injury, astrocytes can form glial scars to seal off injured areas and protect from further damage. Here, we asked if astrocyte loss and opening of the BBB is sufficient to induce a scar-forming response in astrocytes adjacent to ablated regions. We first tested for the phosphorylation of the cytokine and growth factor signal transducer and activator of transcription 3 (pSTAT3), which is essential for scar formation (Herrmann et al., 2008). The promoter for glial fibrillary acidic protein (GFAP) is a downstream target of STAT3 (Ito et al., 2016). GFAP is typically upregulated after injury and is intensely increased in scar-forming astrocytes.

While we found GFAP^+^ astrocytes adjacent to regions of astrocyte ablation and Cadaverine leakage at 1 dpa, 3 dpa, and 5 dpa, at these timepoints none labeled for pSTAT3. Several pockets of pSTAT3^+^ /GFAP^+^ astrocytes were detected at 11 dpa. By 28 dpa, pSTAT3^+^ /GFAP^+^ astrocytes were more numerous and appeared to surround areas of ablation **(Fig. 4a)**. These astrocytes had increased GFAP levels when compared to those at earlier timepoints. These regions were observed across the cortex and were found in all experimental mice at 28 dpa (n=3).

**Figure 4.**
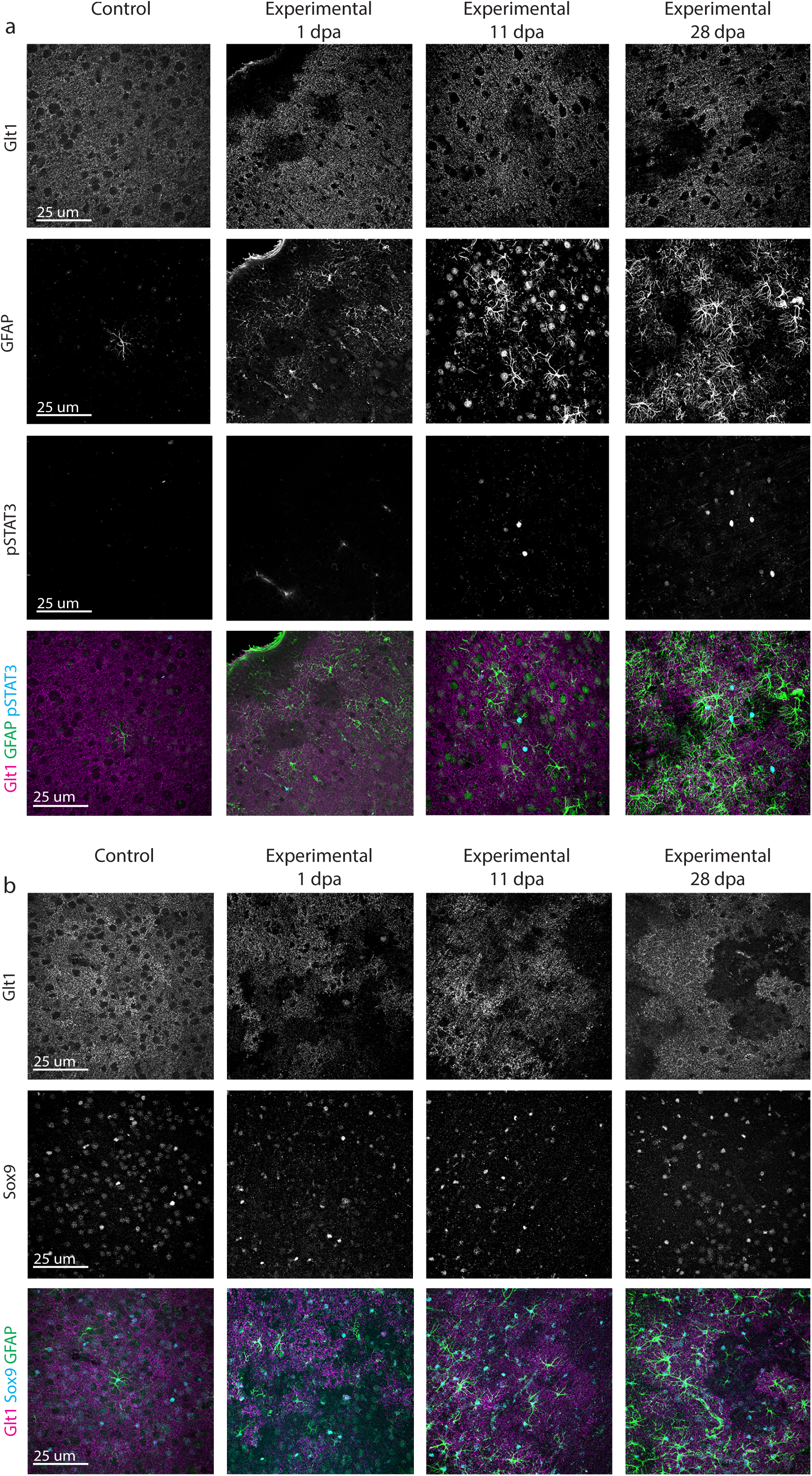
Astrocytes adjacent to areas of astrocyte ablation had late increased phosphorylation of STAT3 and unchanged Sox9. **a.** Astrocytes adjacent to regions of Glt1 loss were negative for pSTAT3 at 1, 3 and 5 dpa. GFAP^+^ astrocytes close to astrocyte ablation areas labeled positive for phosphorylated STAT3 at 11 dpa and 28 dpa. **b.** Astrocytes adjacent to regions of astrocyte ablation indicated by lack of Glt1 did not show changes in their expression of the astrocyte transcription factor Sox9.

We next assessed if astrocyte ablation activated microglia and if this might affect timing of astrocytic STAT3 phosphorylation. At 3 dpa, some microglia had increased levels of Iba1 appeared changed in morphology, with many membrane-rich processes. At 11dpa, many more microglia within areas of astrocyte ablation were clearly activated with Iba1 upregulation, hypertrophy of cell bodies and main processes. At this timepoint, classic morphological changes reminiscent of microglia activation dominated while “membrane-rich” microglia were no longer observed. At 28dpa, microglia activation within the areas of astrocyte ablation was even more pronounced. Some microglia were de-ramified while others had an increased number of processes (**Suppl. Fig. 5**). Given that phosphorylation of STAT3 in astrocytes aligned in severity with the severity of microglia activation is possible that astrocytes and microglia close to ablation areas respond to each other’s activation state.

The transcription factor Sox9 is expressed primarily by astrocytes and is upregulated in reactive astrocytes in models of stroke and Parkinson’s disease (Choi et al., 2018; Sun et al., 2017). We tested for Sox9 expression in astrocytes adjacent to regions of ablation but found that Sox9 expression remained unchanged in all groups **(Fig. 4b)**.

### 3.5 Astrocytes adjacent to ablated areas respond with morphological changes

After focal brain injury, astrocytes become hypertrophic with swollen GFAP^+^ cell bodies and processes. Astrocytes extend their processes toward the site of injury (Oberheim, Wang, Goldman, & Nedergaard, 2006; Robel, Bardehle, Lepier, Brakebusch, & Götz, 2011). This results, in conjunction with the response of other glia, in a physical boundary sealing the injured area off from the uninjured parts of the brain. To determine if astrocytes adjacent to ablated astrocytes respond in a similar fashion, we employed the Simple Neurite Tracer in Fiji/ImageJ in order to create a 3D projection of GFAP^+^ astrocytes and their processes. Length and thickness of astrocytes’ major GFAP-expressing processes were traced, and Sholl analysis was used to determine process complexity. In control mice, sparse numbers of GFAP^+^ astrocytes were located along blood vessels. These astrocytes were traced as controls. After astrocyte ablation, astrocytes neighboring the ablated regions changed over time in the number, length and volume of their GFAP^+^ processes **(Fig. 5a).** Total process length of neighboring astrocytes increased by 43.4 percent from 1 dpa (815.919 µm ± 77.147 µm, n=15) to 28 dpa (1170.540 ± 116.179 µm, n=15) **(Fig. 5b)**. Total process volume of these astrocytes more than doubled, increasing by 140.0 percent from 1 dpa (928.828 µm ± 110.830 µm^3^, n=15) to 28 dpa (2229.636 ± 110.830 µm^3^ n=15) **(Fig. c).** Sholl analysis comparing astrocytes neighboring ablation regions at 1 dpa and 28 dpa showed a significant increase in process arborization (Two-way ANOVA for number of intersections versus timepoint, p<0.0001, n=15 astrocytes per timepoint) **(Fig. 5d),** suggesting that more astrocyte processes expressed GFAP over time.

To determine if astrocytes extend processes into the areas of astrocyte ablation, we examined the dimensions of astrocyte domains by drawing quadrants, using the location of the ablation region as a guide **(Fig. 5e).** We did not find changes in length to width ratio at any timepoint compared to controls (**Fig. 5f).** When looking at the area of the quadrants, we also found no changes in proximal to distal area at any timepoint compared to controls **(Fig. 5g).** This suggests that longer GFAP^+^ processes result from GFAP-filaments stretching farther into already present processes rather than processes stretching into areas of loss to replace lost astrocytes. Alternatively, our approach to measuring astrocyte shape based on GFAP-tracing might not be sensitive enough to detect small polarizations toward areas of loss.

In all, these data suggest that loss of astrocytes initiates mild morphological changes in their neighbors characterized by presence of GFAP filaments in a larger number of processes and increase in process volume, yet we could not detect directional polarity to these changes. Thus, loss of contact to a neighbor might either not be a signal inducing astrocyte polarity or diffuse BBB leakage, other ablation regions in close proximity, and additional cellular and molecular triggers may be present that overwrite a strong polarization toward the lost astrocyte.

### 3.6 Astrocytes adjacent to ablated areas do not proliferate to replace ablated neighbors

Astrocytes in the healthy mature brain are post-mitotic, but a subset of astrocytes re-enter the cell cycle after focal brain or spinal cord injury. We next asked if astrocyte ablation triggers a proliferative response in neighboring astrocytes geared toward replacing lost cells. In order to assess astrocyte proliferation, we chose two approaches. First, we labeled brain slices of experimental and control mice with Ki67, which is expressed during active phases of the cell cycle but is absent in resting non-mitotic cells (Scholzen & Gerdes, 2000). As expected, a small number of Ki67^+^ cells were present throughout the cortical gray matter and a higher number was found within adult neurogenic zones in all groups (**Suppl. Fig. 6a)**. However, S100β^+^ astrocytes adjacent to ablated astrocytes did not co-label with Ki67 within the cortical gray matter at 6 hpa, 1, 3, 5, 11 or 28 dpa (**Suppl. Fig. 6b**, control: 2 Ki67^+^/S100β^+^ cells among 10,088 S100β^+^ cells, n=10 mice; experimental: 1 Ki67/S100β^+^ cell among 3,684 S100β^+^ cells, n= 3-5 mice per timepoint)

Secondly, we labeled all cells that proliferated using the base analog Bromodeoxyuridine (BrdU), which was administered twice daily after Tx administration for 1, 3 and 5 days to cumulatively label all cells that proliferated during this time. BrdU is incorporated during S-phase and can later be detected by immunohistochemistry, even if the cells were not actively proliferating at the time of the tissue harvest. Similar to the findings using the acute proliferation marker Ki67, we did not find S100β^+^ astrocytes that co-labeled for BrdU at 1, 3, or 5 dpa (**Suppl. Fig. 7,** control: 1 BrdU^+^/S100β^+^ cell among 1319 S100β^+^ cells, n=9 mice; experimental: 1 BrdU^+^/S100β^+^ cell among 2865 S100β^+^ cells, n=3 mice per timepoint). Together, astrocyte loss did not initiate proliferation of adjacent astrocytes early or late after ablation.

## 4. Discussion

### 4.1 Astrocyte loss in the adult brain causes BBB damage of variable extent

Here, we demonstrate a necessary and non-redundant role for astrocytes in BBB maintenance *in vivo* using a mouse model of astrocyte ablation via tamoxifen-inducible conditional expression of the non-toxic DTA subunit, which inhibits protein synthesis. Astrocyte apoptosis occurred as early as 2 hours after tamoxifen administration. Cadaverine (∼900 Da) leakage indicative of damage to the BBB occurred just as early around ablated astrocytes. In addition to penetration of the smaller molecular weight Cadaverine into the brain parenchyma, deposition of fibrinogen (340kDa) had taken place in a small subset of the areas with Cadaverine leakage. This suggests varying extents of BBB damage, possibly due to differing numbers of ablated astrocytes. Three recent studies ablating astrocytes have suggested that astrocytes are not needed for maintenance of the BBB *in vivo*. In the spinal cord, astrocyte ablation using a similar genetic DTA ablation system, no obvious lack of BBB integrity was observed (Schreiner et al., 2015; Tsai et al., 2012). Differences in CNS region (cortical gray matter versus spinal cord) or in the genetic approach (Glast-CreERT versus GFAP-CreERT2 or Aldh1l1-loxP-eGFP-STOP-loxP-DTA x Pax3-Cre) could account for the differences. Additionally, these studies only tested for large size damage to the BBB including presence of erythrocytes, lymphocytes and fibrinogen. It would be easy to overlook the small number of areas with BBB damage large enough to permit entry of fibrinogen or cells from the periphery. In our hands, detection of the larger-size breaches required screening of entire brains. The aforementioned studies did not assess smaller size BBB damage. Yet, small openings of the BBB permitting markers below 3 kDa still indicate significant dysfunction in the BBB (Yanagida et al., 2017).

One study concluded that there were no changes to the BBB based on the absence of leakage of Evans Blue (960 Da when unbound and 65-70 kDa when bound to albumin) and dextran (4 kDa) after laser ablation of astrocytes or endfeet followed by 2-photon imaging (Kubotera et al., 2019). Yet cell death was only confirmed by lack of a fluorescent eGFP signal, which can occur as a result of photobleaching after which the still-alive cell is no longer visible. The authors observed recovery of the GFP signal on vessels over the course of minutes to hours but no consequence of the intervention even though free Evans Blue is similar in size as Cadaverine. In our experience, GFP-labeled astrocytes photobleach readily while it is difficult to fully ablate the entire astrocyte or parts of it. It is also possible that 2-photon *in vivo* imaging is not sensitive enough to detect leakage of small amounts of dye. The advantage of Cadaverine is its uptake into neurons, which makes detection of even small leakages easy in tissue processed for immunohistochemistry when compared to detection of diffuse fluorescence in 2-photon imaging. Alternatively, it is possible that ablation of a single astrocyte or single endfoot is not sufficient to induce BBB damage and that secreted factors by neighboring cells are sufficient to maintain barrier integrity. Integrity of tight junctions and other barrier properties were not assessed in this study (Kubotera et al., 2019).

### 4.2 Do astrocyte-secreted factors maintain the blood-brain barrier?

We tested for tight junction integrity by examining the essential tight junction protein ZO-1, a transmembrane protein needed for assembly of the tight junction complex. In order to anchor the tight junctional complex to the cell’s actin cytoskeleton it must be localized correctly (Katsuno et al., 2008) and any abnormal localization of ZO-1 is indicative of defective tight junctions. Endothelial tight junctions were impaired hours after astrocyte ablation. In areas where astrocytes were ablated but Cadaverine leakage had not yet occurred, tight junctions appeared intact suggesting that astrocyte ablation caused damage to tight junctions, which then resulted in dye leakage. The mechanisms by which astrocytes maintain the BBB need to be resolved in future studies.

Of the secreted factors reported to modulate BBB function, only angiotensinogen is highly expressed in mature astrocytes. Angiotensinogen found in the CNS is produced primarily by astrocytes (Stornetta, Hawelu-Johnson, Guyenet, & Lynch, 1988) and is an essential component of the brain renin-angiotensin system (RAS) that maintains blood pressure homeostasis(Tanimoto et al., 1994).The enzymatic cleavage of angiotensinogen produces angiotensin-II (Ang-II), which binds to BBB endothelial cells and is considered to be a central player in regulating blood pressure, yet there is significant controversy whether Ang-II promotes or disrupts BBB integrity. Studies performed in vitro have shown that Ang-II reduces BBB permeability and promotes tight junction expression in developing BBB endothelial cell monolayers (Wosik et al., 2007), but increases permeability when added to already established monolayers (Fleegal-DeMotta, Doghu, & Banks, 2009). BBB-disruptive effects of Ang-II were also found in vivo, though in the context of injury, disease (Takane et al., 2017), or pre-induced hypertension. In these models, Ang-II was introduced either directly into the CNS via infusion (Li et al., 2016) or indirectly via BBB disruption (Biancardi, Son, Ahmadi, Filosa, & Stern, 2014)that permitted entry of Ang-II (alongside other blood-borne factors) from circulation. Both scenarios resulted in higher than physiological-levels of Ang-II that are not solely derived from astrocytes, and are thus insufficient models for studying roles of Ang-II mediated astrocyte maintenance of the BBB.

Complete lack of angiotensinogen and thus Ang-II also results in BBB disruption, evidenced by diffuse BBB leakage and decreased tight junction expression found in angiotensinogen knockout mice (Kakinuma et al., 1998; Wosik et al., 2007). Although tight junctions were not assessed thoroughly in this study, the BBB dysfunction bears similarities to our data. However, the lack of angiotensinogen throughout the body and during development creates a pre-existing hypertensive environment (Tanimoto et al., 1994) that prevents definitive conclusions about the role of angiotensinogen/ Ang-II in maintenance of the healthy adult BBB.

### 4.3 Astrocyte loss triggers partial and delayed scar formation in neighboring astrocytes

In areas surrounding astrocyte ablation, neighboring astrocytes responded with mild to moderate GFAP increase within the first week after ablation. This occurred in conjunction with microglia activation, which occurred around 3 days after ablation and was pronounced in areas of astrocyte loss at 28 dpa. Phosphorylation of the transcriptional activator STAT3, which has a binding site within the GFAP promoter and occurs 1-5 days after spinal cord injury, is responsible for scar formation after spinal cord injury (Wanner et al., 2013). Interestingly, STAT3 phosphorylation did not occur until 11 days after astrocyte ablation, well after we noted an increase in GFAP. This is in contrast to other reports demonstrating that phosphorylation of STAT3 precedes GFAP upregulation in striatal neurotoxicity (O’Callaghan, Kelly, VanGilder, Sofroniew, & Miller, 2014) and after spinal cord injury. In the latter injury paradigm, STAT3 phosphorylation subsides 7-14 days later (Herrmann et al., 2008). However, 2-4 weeks after astrocyte ablation an increasing number of astrocytes were positive for pSTAT3 which correlated with increased numbers of GFAP^+^ processes and morphological changes including cellular hypertrophy. This delayed timeline for STAT3 phosphorylation might be explained by the gradual astrocyte loss and reduced extent of BBB damage, compared to an acute large invasive injury with hemorrhage and infiltration of immune cells. Whether astrocyte cell death or BBB leakage triggered the response of neighboring astrocytes was difficult to determine given that BBB damage occurred quickly after astrocyte ablation.

### 4.4 Astrocyte loss does not induce proliferation of neighboring astrocytes

Astrocytes adjacent to ablated areas did not re-enter the cell cycle suggesting that loss of a neighbor is not sufficient to trigger re-entry into the cell cycle. This is surprising as culture studies suggest that contact inhibition is a prominent signal suppressing cell proliferation and exit from the cell cycle (Pavel et al., 2018). The lack of proliferation after astrocyte ablation in the adult brain suggests that other signals trigger re-entry of astrocytes into the cell cycle after CNS injury. In the context of severe focal brain or spinal cord injury, a 30-70 percent of scar-forming astrocytes proliferate (Bardehle et al., 2013; Buffo et al., 2008; Wanner et al., 2013), while proliferation is not initiated after genetically induced neuronal cell death, in mouse models of Alzheimer disease (Behrendt et al., 2013), or in a mouse model of chronic astrogliosis (Robel et al., 2009). Based on these observations, exposure of the brain to blood-borne substances might be a possible trigger initiating proliferation after acute injury. However, we did not observe proliferating astrocytes in areas with Cadaverine or fibrinogen leakage. This does not exclude the possibility that rapid exposure to or higher concentrations of blood-borne factors would induce astrocyte proliferation. Alternatively, different factors or a combination of factors absent after astrocyte ablation are necessary for astrocytes to re-enter the cell cycle.

### 4.5 The blood-brain barrier fails to repair in areas of astrocyte ablation

We observed BBB damage across all timepoints, suggesting a lack of barrier repair. However, the area covered by Cadaverine leakage was reduced by half and appeared restricted to areas of astrocyte ablation at 28 dpa, suggesting that the response of adjacent astrocytes might restrict diffusion of blood-borne factors. In the context of CNS injury, a dual role for astrocytes at the BBB is documented by many studies: In some injury paradigms, astrocytes upregulate factors that increase BBB permeability via downregulation of tight junctions. Among these factors are vascular endothelial growth factor, matrix metalloproteinases, and nitric oxides (Gu et al., 2012; Jiang, Xia, Jiang, Wang, & Gao, 2014; Yang, Estrada, Thompson, Liu, & Rosenberg, 2007). Conversely, increased expression of glial derived neurotrophic factor and retinoic acid promote tight junction expression and reduce permeability after stroke (Kong et al., 2015; Liu et al., 2016) and in multiple sclerosis (Mizee et al., 2014). Lack of angiotensinogen has also been associated with failure of BBB repair after cold injury. While the BBB was repaired and not permissive to Evans Blue in wildtype mice within 5 days of the injury, Evans Blue still leaked into the brain parenchyma of angiotensinogen KO mice 2 weeks post injury (Kakinuma et al., 1998). Thus dependent on disease context, age and surrounding microenvironment, astrocytes may lose their ability to maintain the BBB, contribute to BBB repair or even cause barrier damage.

## Acknowledgements

This work was supported by the National Institute of Neurological Disorders and Stroke at the National Institutes of Health (grant number R01NS105807). We thank Katie Barnes, Maame Boateng, Deyton Cook, and Dzenis Mahmutovic for their contributions to data collection.

## Conflict of Interest Statement

The authors declare no conflict of interest.

## Data Availability Statement

The data supporting the findings in this study are available upon reasonable request from the corresponding author.

---

**Supplemental Figure 1.**
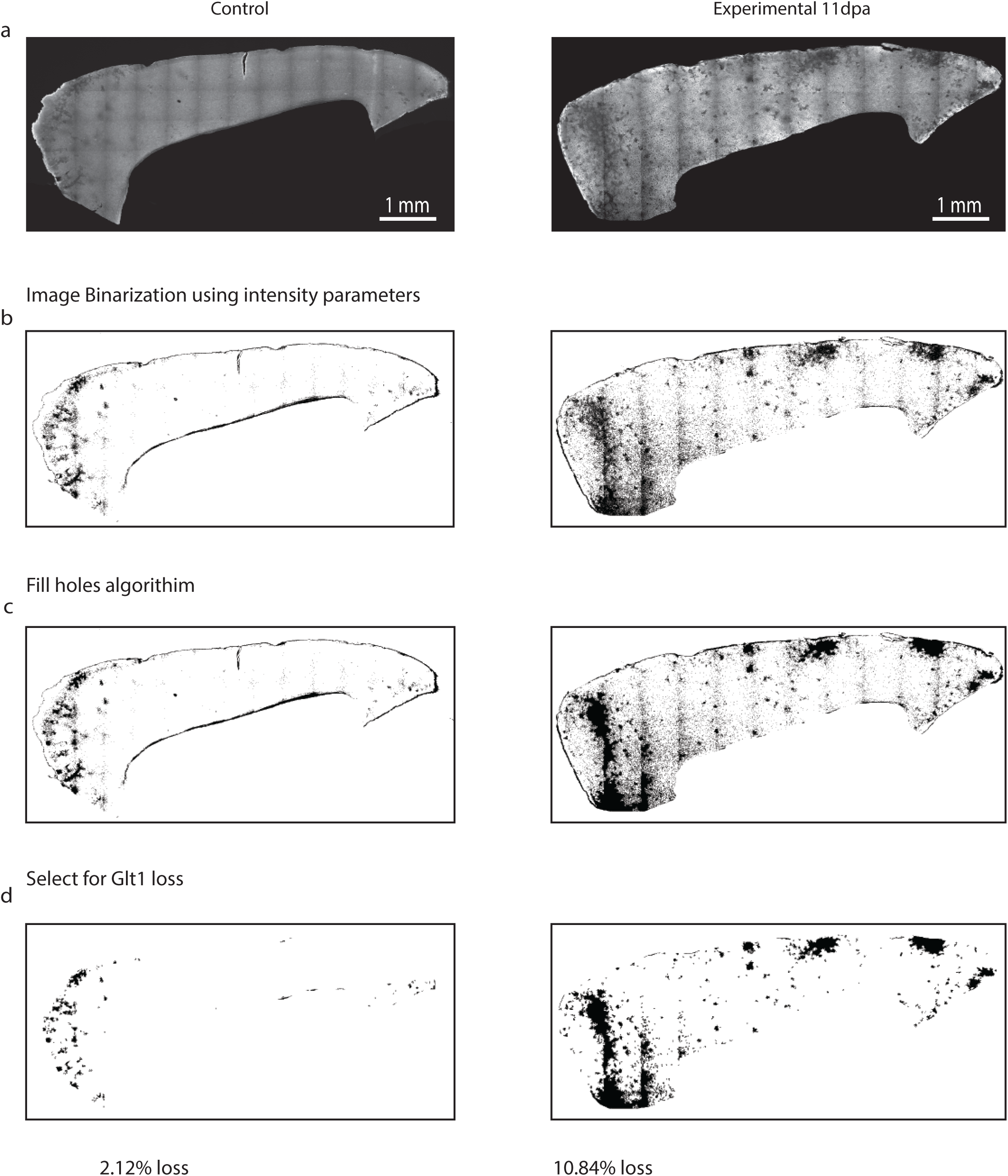
Semi-automatic quantification of Glt1 loss regions using ImageJ. **a.** Large images of sagittal slices were acquired for both experimental and control mice at all timepoints using line scanning at 20x magnification. **b.** Images were binarized based using intensity parameters that selected for Glt-1 negative regions. **c.** Binary images were run through the “fill holes” algorithm to fill in Glt-1 negative regions and create discrete shapes. **d.** The Glt1 negative regions that are blood vessels and neurons were excluded using consistent size and circularity quantification parameters. The remaining Glt1 negative regions were quantified as regions of astrocyte ablation.

**Supplemental Figure 2.**
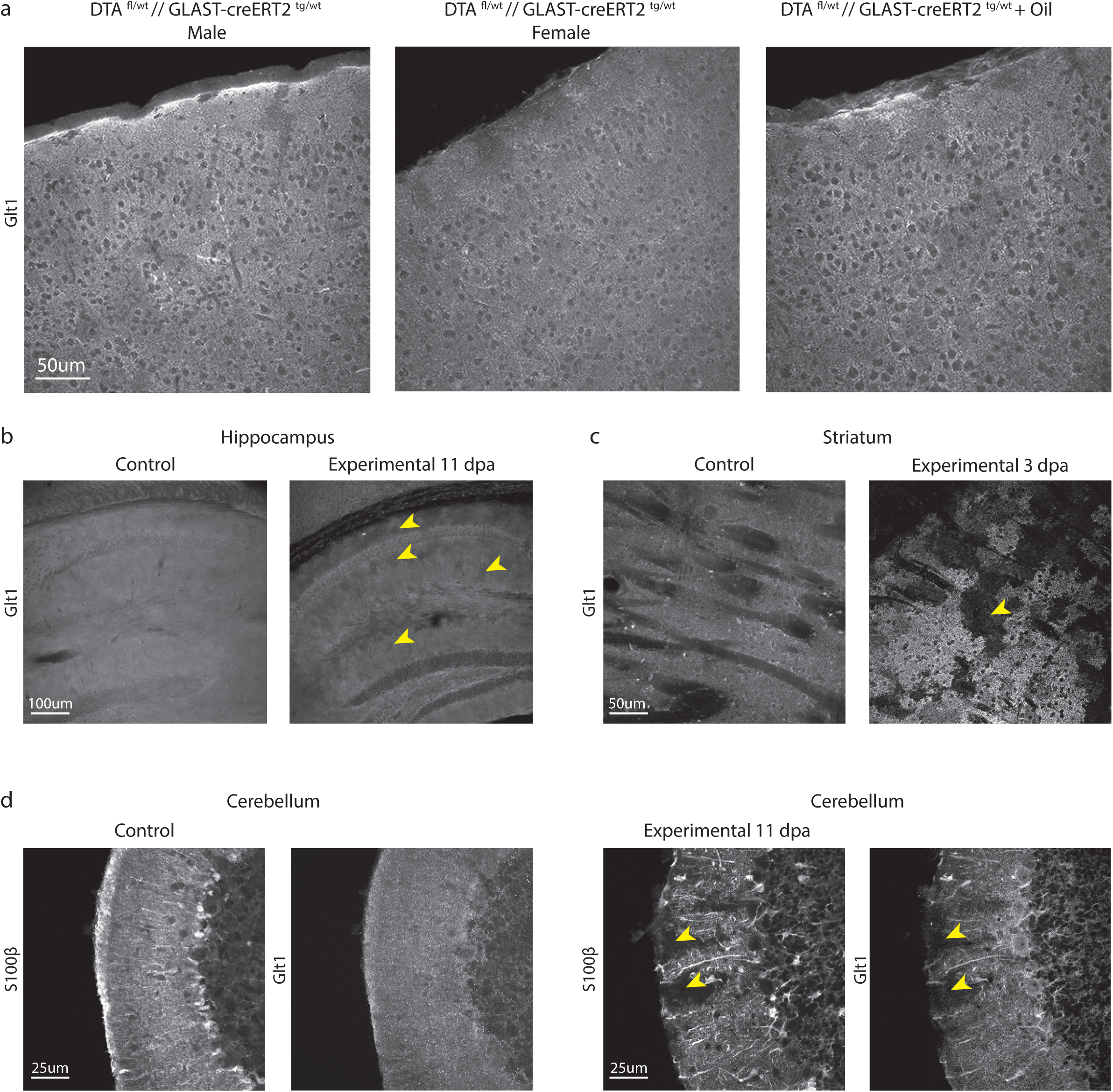
a. Male and female mice (DTA^fl/wt^//Glast-CreERT^tg/wt^) administered tamoxifen or an oil vehicle only or nothing at all (naïve) did not show astrocyte ablation indicated by no lack of the continuous labeling of astrocyte fine processes with the astrocyte marker membrane-associated astrocyte glutamate transporter Glt1. **b-c.** In addition to cortical loss of Glt1, Experimental mice showed Glt1 loss in hippocampus and striatum after tamoxifen administration. **d.** Bergmann glia in the cerebellum that highly express GLAST at high levels in the adult and this in cerebellum showed noticeable depletion in density after tamoxifen administration (arrowheads).]

**Supplemental Figure 3.**
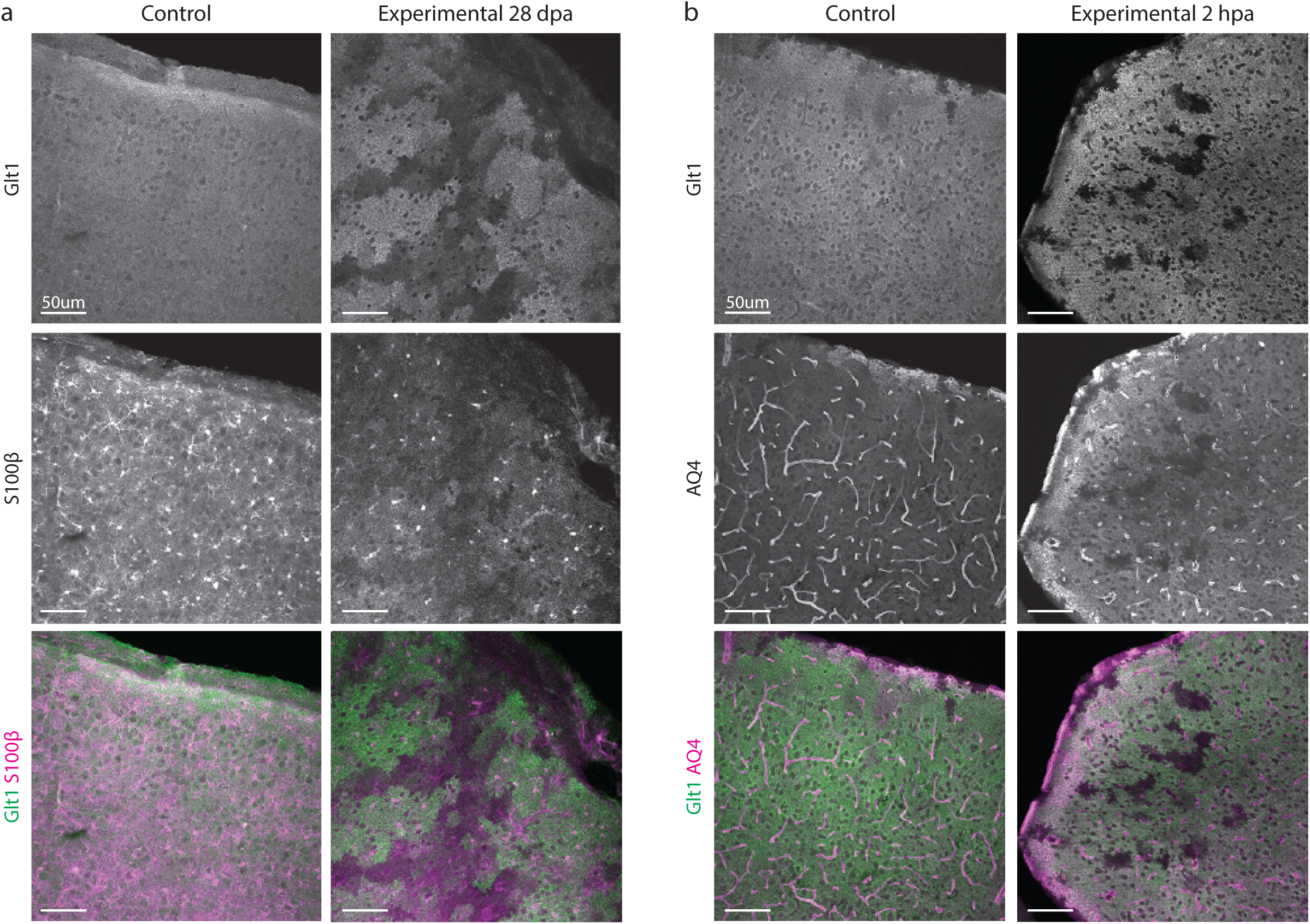
Astrocyte ablation in cortex indicated by lack of expression of astrocyte markers. **a.** Lack of astrocyte marker S100β overlaps with lack of Glt1 in experimental mice. **b.** Lack of astrocyte marker Aquaporin-4 (AQ4) overlaps with lack of Glt1 in experimental mice.

**Supplemental Figure 4.**
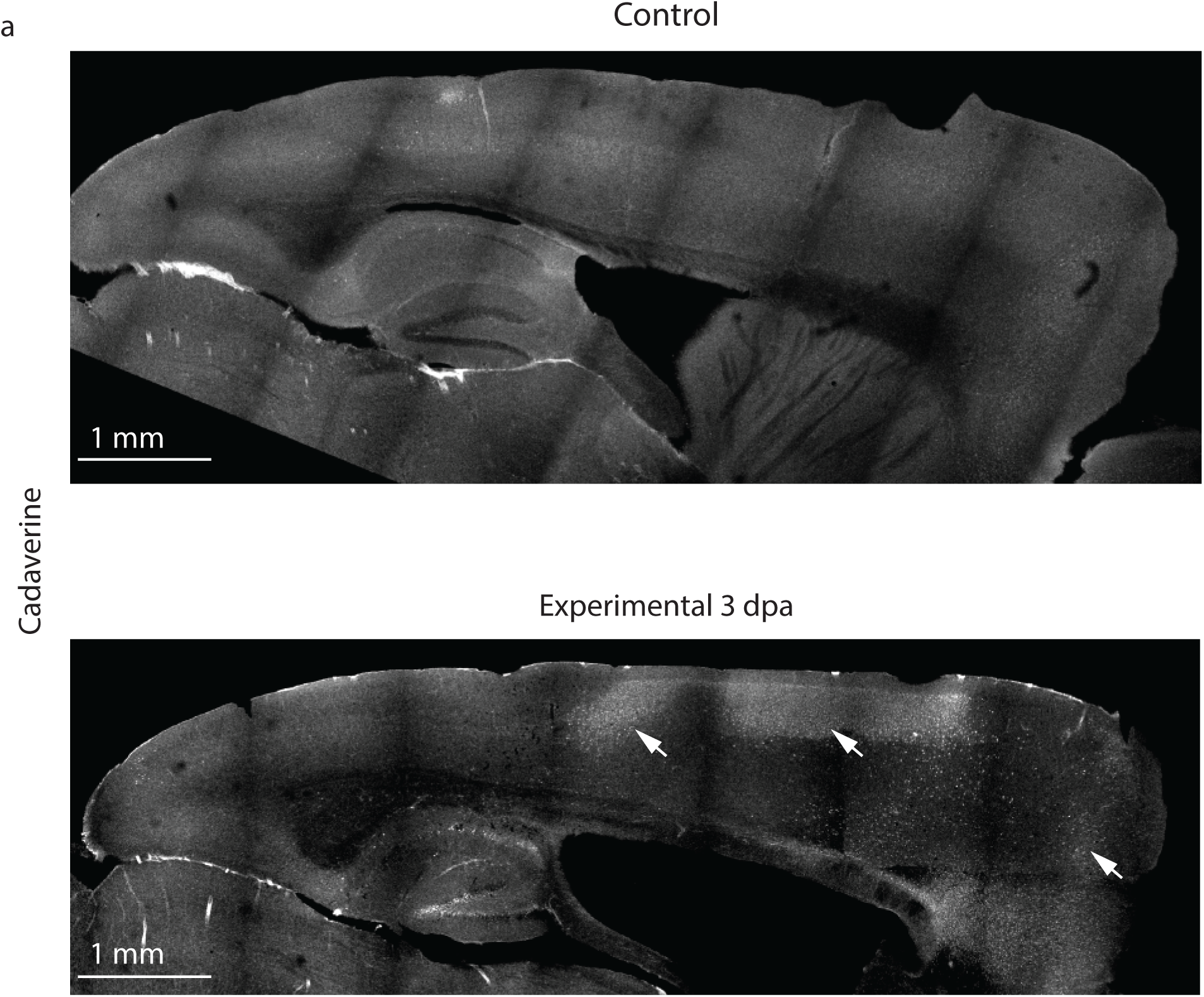
Cadaverine leakage occurred after astrocyte ablation across cortex. **a.** Large area scans of cortex showed minimal Cadaverine leakage in control mice, while diffuse areas of leakage were found in experimental mice.

**Supplemental Figure 5.**
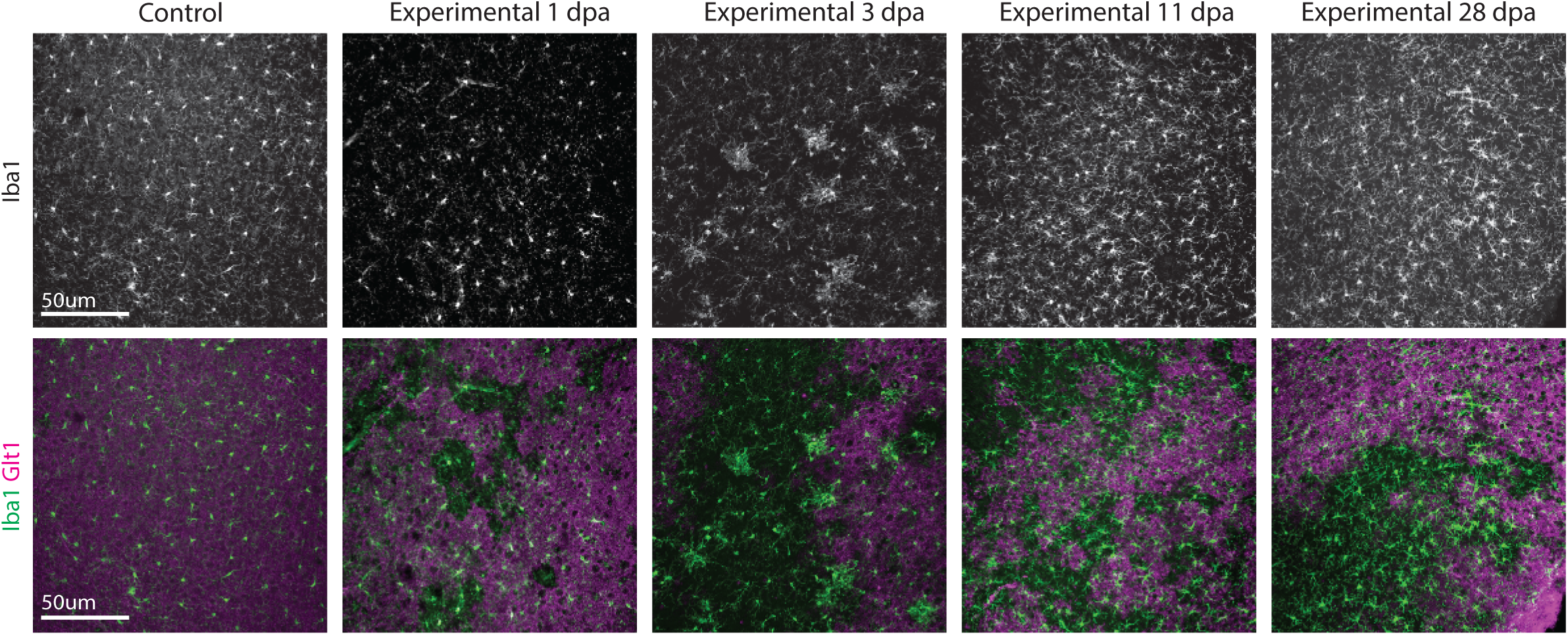
Microglia showed morphology changes and increased Iba1 levels in regions of ablated astrocytes.

**Supplemental Figure 6.**
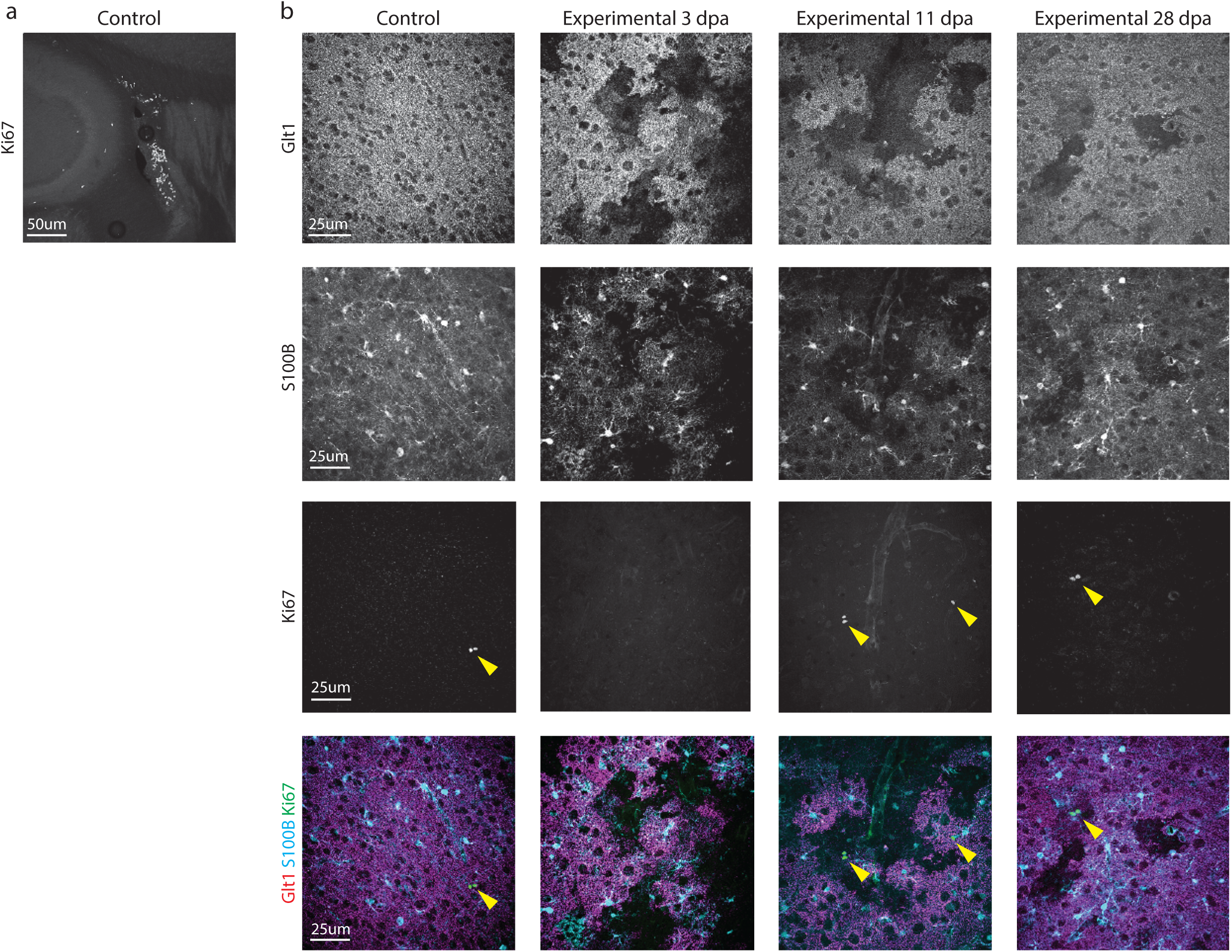
Astrocyte ablation does not induce proliferation of neighboring astrocytes. **a.** Ki67^+^cells were present in all mice at the subependymal zone, which contains proliferating cells in adult mice. **b.** Astrocytes adjacent to loss regions did not express the mitotic marker Ki67 but Ki67^+^ cells of other cell identity (arrowheads) were present in both control and experimental groups.

**Supplemental Figure 7.**
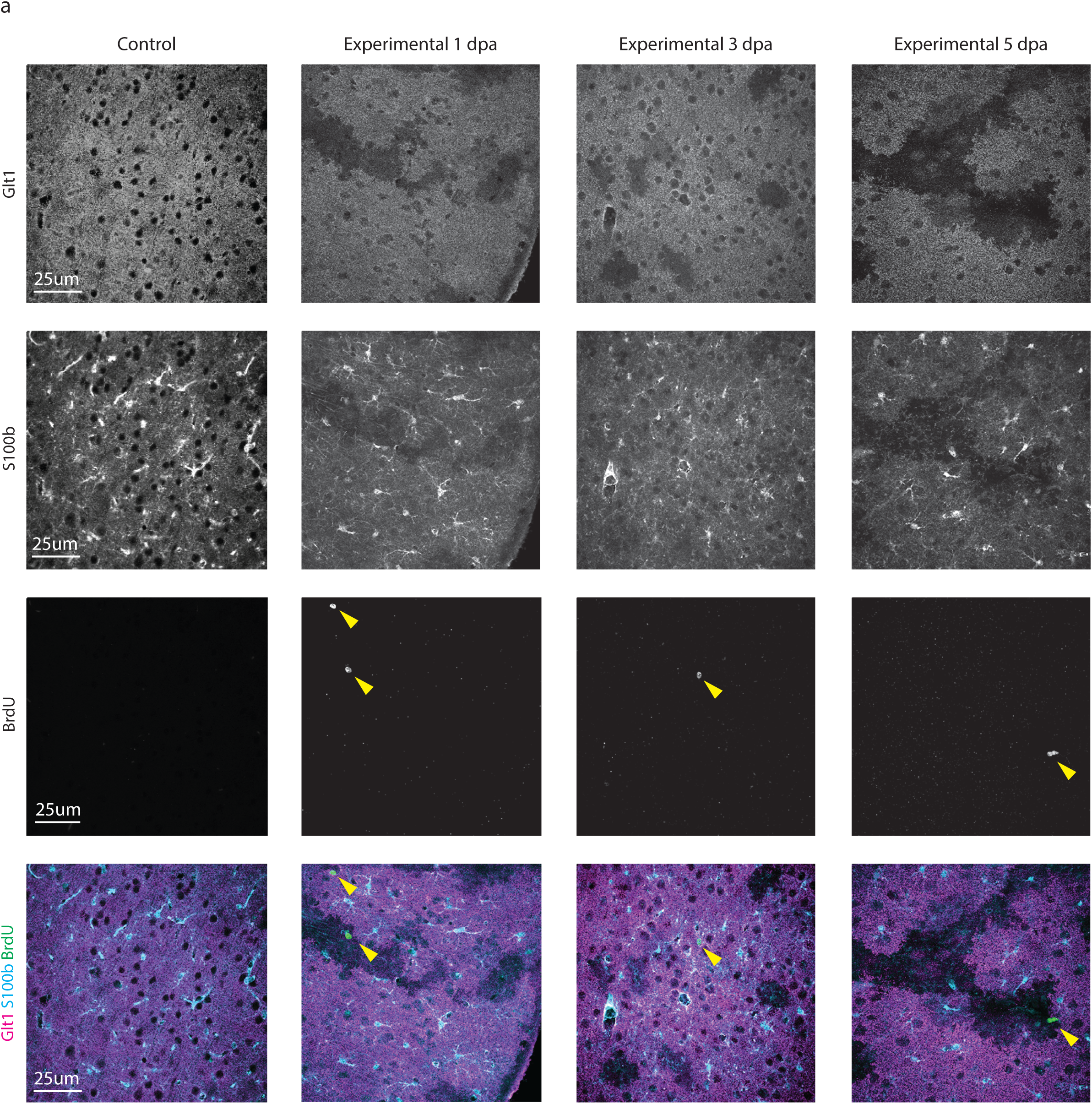
Astrocyte ablation does not induce proliferation of neighboring astrocytes. **a.** Astrocytes adjacent to loss regions did not express the mitotic marker BrdU but BrdU^+^ cells of other cell identity (arrowheads) were present in both control and experimental groups.

